# A novel membrane protein Hoka regulates septate junction organization and stem cell homeostasis in the *Drosophila* gut

**DOI:** 10.1101/2020.11.10.376210

**Authors:** Yasushi Izumi, Kyoko Furuse, Mikio Furuse

## Abstract

Smooth septate junctions (sSJs) regulate the paracellular transport in the intestinal and renal system in arthropods. In *Drosophila*, the organization and physiological function of sSJs are regulated by at least three sSJ-specific membrane proteins: Ssk, Mesh, and Tsp2A. Here, we report a novel sSJ membrane protein Hoka, which has a single membrane-spanning segment with a short extracellular region having 13-amino acids, and a cytoplasmic region with three repeats of the Tyr-Thr-Pro-Ala motif. The larval midgut in *hoka*-mutants shows a defect in sSJ structure. Hoka forms a complex with Ssk, Mesh, and Tsp2A and is required for the correct localization of these proteins to sSJs. Knockdown of *hoka* in the adult midgut leads to intestinal barrier dysfunction, stem cell overproliferation, and epithelial tumors. In *hoka*-knockdown midguts, aPKC is up-regulated in the cytoplasm and the apical membrane of epithelial cells. The depletion of *aPKC* and *yki* in *hoka*-knockdown midguts results in reduced stem cell overproliferation. These findings indicate that Hoka cooperates with the sSJ-proteins Ssk, Mesh, and Tsp2A to organize sSJs, and is required for maintaining intestinal stem cell homeostasis through the regulation of aPKC and Yki activities in the *Drosophila* midgut.

**Summary statement:** Depletion of *hoka*, a gene encoding a novel septate junction protein, from the *Drosophila* midgut results in the disruption of septate junctions, intestinal barrier dysfunction, stem cell overproliferation, and epithelial tumors.

## Introduction

Epithelia separate distinct fluid compartments within the bodies of metazoans. For this epithelial function, occluding junctions act as barriers that control the free diffusion of solutes through the paracellular pathway. Septate junctions (SJs) are occluding junctions in invertebrates (Furuse and Tsukita, 2006; Lane, 1994; Tepass and Hartenstein, 1994) and form circumferential belts along the apicolateral region of epithelial cells. In transmission electron microscopy, SJs are observed between the parallel plasma membranes of adjacent cells, with ladder-like septa spanning the intermembrane space (Lane, 1994, Tepass and Hartenstein, 1994). Arthropods have two types of SJs: pleated SJs (pSJs) and smooth SJs (sSJs) (Banerjee et al., 2006; Lane, 1994; Tepass and Hartenstein, 1994; Jonusaite et al., 2016). pSJs are found in ectoderm derived epithelia and surface glia surrounding the nerve cord, whereas sSJs are found mainly in the endoderm derived epithelia, such as the midgut and gastric caeca (Lane, 1994; Tepass and Hartenstein, 1994). Despite being derived from the ectoderm, the outer epithelial layer of the proventriculus (OELP) and the Malpighian tubules also possess sSJs (Lane, 1994; Tepass and Hartenstein, 1994). Further, pSJs and sSJs are distinguished by the arrangement of septa. For example, the septa of pSJs form regular undulating rows, whereas those in sSJs form regularly spaced parallel lines in the oblique sections in lanthanum-treated preparations (Lane and Swales, 1982; Lane, 1994). To date, more than 20 pSJ-related proteins have been identified and characterized in *Drosophila* (Banerjee et al., 2006; Izumi and Furuse, 2014; Tepass et al., 2001; Wu and Beitel, 2004; Rouka et al., 2020). In contrast, only three membrane-spanning proteins, i.e., Ssk, Mesh, and Tsp2A have been reported as specific molecular constituents of sSJs (sSJ-proteins) in *Drosophila* (Furuse and Izumi, 2017). Therefore, the mechanisms underlying sSJ organization and functional properties of sSJs remain poorly understood compared with pSJs. Ssk has four membrane-spanning domains; two short extracellular loops, cytoplasmic N- and C-terminal domains, and a cytoplasmic loop (Yanagihashi et al., 2012). Mesh is a cell-cell adhesion molecule, which has a single-pass transmembrane domain and a large extracellular region containing a NIDO domain, an Ig-like E set domain, an AMOP domain, a vWD domain, and a sushi domain (Izumi et al., 2012). Tsp2A is a member of the tetraspanin family of integral membrane proteins in metazoans with four transmembrane domains, N- and C-terminal short intracellular domains, two extracellular loops, and one short intracellular turn (Izumi et al., 2016). The loss of *ssk*, *mesh* and *Tsp2A* causes defects in the ultrastructure of sSJs and the barrier function against a 10 kDa fluorescent tracer in the *Drosophila* larval midgut (Yanagihashi et al., 2012; Izumi et al., 2012; Izumi et al., 2016). Ssk, Mesh, and Tsp2A interact physically and are mutually dependent for their sSJ localization (Izumi et al., 2012; Izumi et al., 2016). Thus, Ssk, Mesh, and Tsp2A act together to regulate the formation and barrier function of sSJs. Further, Ssk, Mesh, and Tsp2A are localized in the epithelial cell-cell contact regions in the *Drosophila* Malpighian tubules where sSJs are present (Tepass and Hartenstein, 1994; Yanagihashi et al., 2012; Izumi et al., 2012; Izumi et al., 2016). Recent studies have shown that the knockdown of *mesh* and *Tsp2A* in the epithelium of Malpighian tubules leads to defects in epithelial morphogenesis, tubule transepithelial fluid and ion transport, and paracellular macromolecule permeability in the tubules (Jonusaite et al., 2020; Beyenbach et al., 2020). Thus, sSJ-proteins are involved in the development and maintenance of functional Malpighian tubules in *Drosophila*.

The *Drosophila* adult midgut consists of a pseudostratified epithelium, which is composed of absorptive enterocytes (ECs), secretory enteroendocrine cells (EEs), intestinal stem cells (ISCs), EC progenitors (enteroblasts: EBs), and EE progenitors (enteroendocrine mother cells: EMCs) (Micchelli and Perrimon, 2006; Ohlstein and Spradling, 2006; Guo and Ohlstein, 2015). The sSJs are formed between adjacent ECs and between ECs and EEs (Resnik-Docampo et al., 2017). To maintain midgut homeostasis, ECs and EEs are continuously renewed by proliferation and differentiation of the ISC lineage through the production of intermediate differentiating cells, EBs and EMCs. Recently, we and other groups reported that the knockdown of sSJ-proteins Ssk, Mesh, and Tsp2A in the midgut causes intestinal hypertrophy accompanied by the overproliferation of ECs and ISC (Salazar et al., 2018; Xu et al., 2019; Izumi et al., 2019, Chen et al., 2020). These results indicate that sSJs play a crucial role in maintaining tissue homeostasis through the regulation of stem cell proliferation and enterocyte behavior in the *Drosophila* adult midgut. Furthermore, Chen et al., (2018) have reported that the loss of *mesh* and *Tsp2A* in adult midgut epithelial cells causes defects in cellular polarization. although no remarkable defects in epithelial polarity were found in the first-instar larval midgut cells of *ssk*, *mesh*, and *Tsp2A*-mutants (Yanagihashi et al., 2012; Izumi et al., 2012; Izumi et al., 2016). Thus, sSJs may contribute to the establishment of epithelial polarity in the adult midgut.

During regeneration of the *Drosophila* adult midgut epithelium, various signaling pathways are involved in the proliferation and differentiation of the ISC lineage (Jiang et al., 2016). Atypical Protein kinase C (aPKC) is an evolutionarily conserved key determinant of apical-basal epithelial polarity (Ohno et al., 2015). Importantly, Chen et al., (2018) have reported that aPKC is dispensable for the establishment of epithelial cell polarity in the *Drosophila* adult midgut. Goulas et al., (2012) have reported that aPKC is required for differentiation of the ISC linage in the midgut. The Hippo signaling pathway is involved in maintaining tissue homeostasis in various organs (Zheng and Pan, 2019). In the *Drosophila* midgut, inhibition of the Hippo signaling pathway activates the transcriptional co-activator Yorkie (Yki), which results in accelerated ISC proliferation via the Unpaired (Upd)-Jak-Stat signaling pathway (Karpowicz et al., 2010; Ren et al., 2010; Shaw et al., 2010). Recent studies have shown that Yki is involved in ISC overproliferation caused by the depletion of sSJ-proteins in the midgut (Xu et al., 2019; Chen et al., 2020). Furthermore, Xu et al. (2019) have shown that aPKC is activated in the *Tsp2A*-RNAi treated midgut, leading to activation of its downstream target Yki and causing ISC overproliferation through the activation of the Upd-JAK-Stat signaling pathway. Thus, crosstalk between aPKC and the Hippo signaling pathways appear to be involved in ISC overproliferation caused by *Tsp2A* depletion.

To further understand the molecular mechanisms underlying sSJ organization, we performed a deficiency screen for Mesh localization and identified the integral membrane protein Hoka as a novel component of *Drosophila* sSJs. Hoka consists of a short extracellular region and the characteristic repeating four-amino acid motifs in the cytoplasmic region, and is required for the organization of sSJ structure in the midgut. Hoka and Ssk, Mesh, and Tsp2A show interdependent localization at sSJs and form a complex with each other. The knockdown of *hoka* in the adult midgut results in intestinal barrier dysfunction, aPKC- and Yki-dependent ISC overproliferation, and epithelial tumors. Thus, Hoka plays an important role in sSJ organization and in maintaining ISC homeostasis in the *Drosophila* midgut.

## Results

### Hoka is involved in sSJ formation

We previously queried *Drosophila* strains that were defective in sSJ accumulation of Mesh using a genetic screen for a chromosomal deficient stock from the Bloomington Deficiency Kit. We found several deficiencies that caused an abrogated sSJ accumulation of Mesh in the stage 16 embryo OELP (Izumi et al., 2016). The *Tsp2A* gene was identified as being responsible for Mesh localization in the screen (Izumi et al., 2016), and we found that the OELP of Df(3L)BSC371 (deleted segment: 64C1-64E1) showed cytoplasmic distribution of Mesh. To identify the precise genomic region responsible for the phenotype, we observed the Mesh distribution with other deficiencies overlapping with Df(3L)BSC371. The OELP of Df(3L)ED210 (deleted segment: 64B9-64C13) (Fig. 1A, B) and Df(3L)Exel6102 (deleted segment: 64B13-64C4) exhibited a cytoplasmic distribution phenotype for Mesh but not that of Df(3L)Exel6103 (deleted segment: 64C4-64C8), and the phenotype for Df(3L)BSC371 was mapped to the 64C1-64C4 interval. Within the genomic region 64C1-64C4, we focused on *CG13704* (Fig. 1C) as it was highly expressed in the midgut and Malpighian tubules (http://flybase.org/reports/FBgn0035583). *CG13704* encodes a putative single-pass transmembrane protein of 136 amino acids with a signal peptide and a transmembrane region (Fig. 1E), which is conserved in insects alone. We named the CG13704 protein ‘Hoka’ based on its immunostaining images in the midgut (see below; Hoka means honeycomb in Japanese). The mature Hoka protein appears to have a short N-terminal extracellular region (13 amino acids) after cleavage of the signal peptide (https://www.uniprot.org/uniprot/Q8SXS4). Interestingly, the C-terminal region is threonine-rich and includes three tyrosine-threonine-proline-alanine (YTPA) motifs. Multiple sequence alignment of Hoka using Multiple Alignment using Fast Fourier Transform (MAFFT) revealed that three types of Hoka homologs (three, two, and single YTPA motif-containing homologs) are present in *Drosophila* (Fig. 1E, S1). The mosquito homologs have a single YTPA motif, and the butterfly homologs have a single similar YQPA motif (Fig. S1).

**Figure 1.**
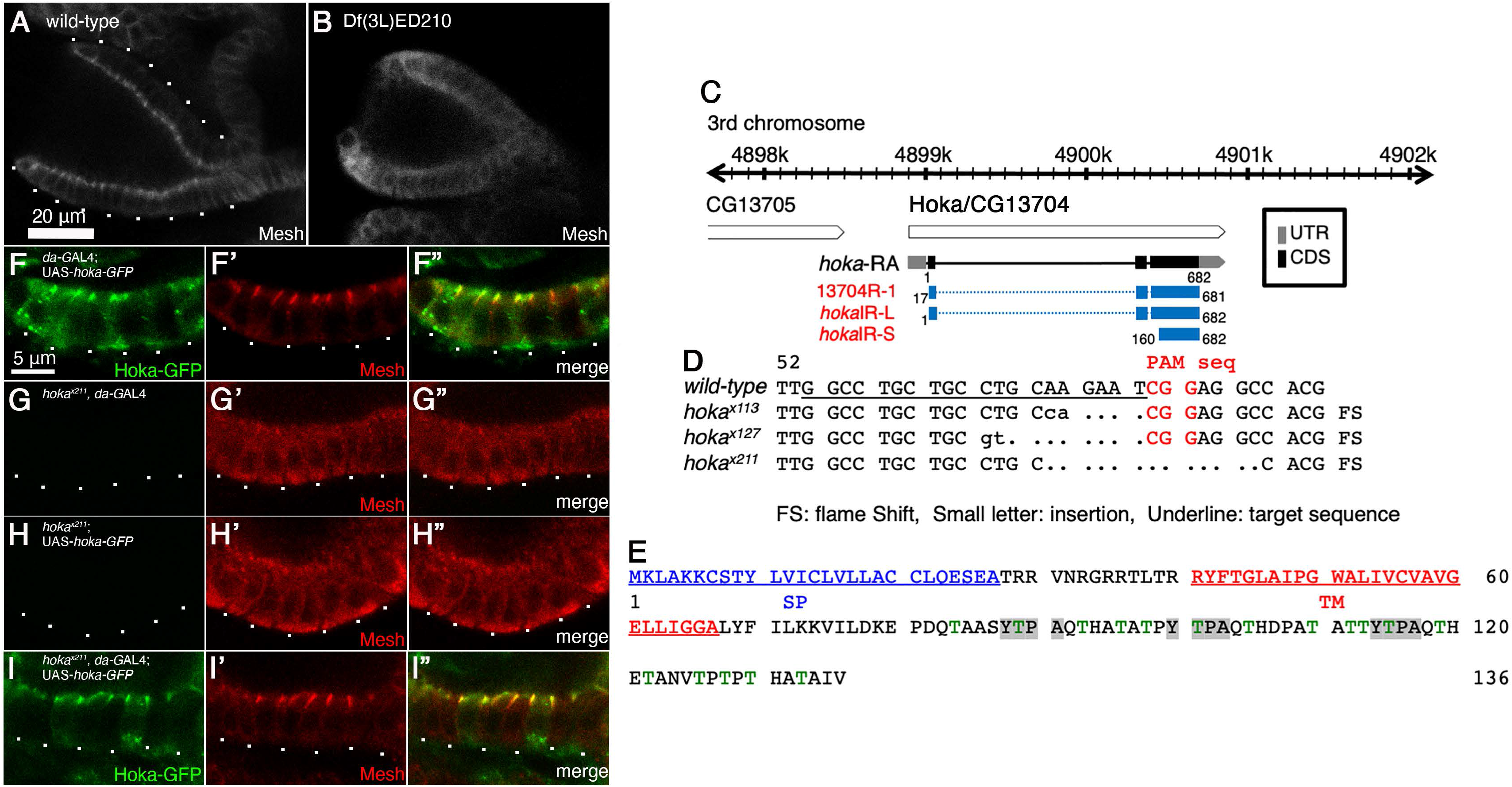
Identification of *hoka* as an sSJ-related gene via a deficiency screen. **(A, B)** Immunofluorescence staining of embryonic stage 16 wild-type (A) and Df(3L)ED210 (B) OELPs using an anti-Mesh antibody. Basal membranes are delineated by dots (A). Scale bar: 20 μm. **(C)** Physical map of the *Drosophila* 3rd-chromosome containing the *hoka* gene. Full-length genes of *hoka/CG13704* are contained in this region. The *hoka* DNA sequence used for the construction of *hoka*-RNAi (13704R-1, *hoka*IR-L, and *hoka*IR-S) is indicated by blue lines. Gray bar: untranslated regions of the *hoka* transcript/*hoka*-RA. Black bar: coding sequences of the *hoka* transcript/*hoka*-RA. **(D)** Genomic sequences of *hoka*-mutations induced by the CRISPR/Cas9 method. The nucleotide sequence of wild-type *hoka* from the start codon is shown at the top. The guide RNA target sequence is underlined and the PAM sequence is shown in red. Deleted nucleotides are shown as dashes. Inserted nucleotides are shown in lowercase letters. **(E)** Amino acid sequence of Hoka. The Hoka polypeptide contains a signal peptide (SP: blue), a transmembrane region (TM: red), and three Tyr-Thr-Pro-Ala (YTPA, highlighted by gray) repeat motifs in the threonine-rich (green) cytoplasmic region. **(F–F”)** Immunofluorescence staining of the Hoka-GFP-expressing stage 16 OELP (*da*-GAL4; UAS-*hoka-GFP*) using anti-GFP (F, F”) and anti-Mesh (F’, F”) antibodies. Hoka-GFP colocalizes with Mesh at sSJs. **(G–I”)** Immunofluorescence staining of *hoka^x211^*-mutant (*hoka^x211^, da*-GAL4, G–G”; *hoka^x211^*, UAS-*hoka-GFP*, H–H”) and Hoka-GFP-expressing *hoka^x211^*-mutant (*hoka^x211^, da*-GAL4, UAS-*hoka-GFP*, I–I”) stage 16 OELPs using anti-GFP (G, G”, H, H”, I, I”) and anti-Mesh (G’, G”, H’, H”, I’, I”) antibodies. Basal membranes are delineated by dots. Scale bar (F–I”): 5 μm.

To examine whether Hoka is associated with sSJs, we expressed C-terminally GFP-tagged Hoka (Hoka-GFP) in flies using *da*-GAL4 (Fig. 1F–F,” see below). In the stage 16 OELP, Hoka-GFP was detected in the apicolateral region with some cytoplasmic aggregates. Mesh colocalized with Hoka-GFP in the apicolateral region (Fig. 1F’, F”), and therefore, we characterized Hoka as an sSJ-associated molecule.

To investigate whether Hoka is involved in sSJ formation, we generated *hoka*-mutants using the CRISPR/Cas9 method provided by NIG-Fly (Kondo and Ueda, 2013). We obtained three independent *hoka*-mutant strains (*hoka^x113^, hoka^x127^*, and *hoka^x211^*), all of which had small indel mutations encompassing the target site (Fig. 1D). These *hoka*-mutant embryos hatched into first-instar larvae but died at this stage (data not shown). All *hoka*-mutants had frameshifts and premature stop codons. In the *hoka^x113^, hoka^x127^*, and *hoka^x211^* mutant stage 16 OELP, Mesh was diffusely distributed in the cytoplasm (Fig. S2F’–I”). Among these mutant strains, we mainly used the *hoka^x211^* mutant for further experiments. To confirm that the lack of *hoka* caused cytoplasmic distribution of Mesh, we expressed Hoka-GFP in *hoka*-mutant flies using *da*-GAL4. The apicolateral accumulation of Mesh in the OELP was recovered and Hoka-GFP colocalized with Mesh in the Hoka-GFP expressing *hoka*-mutant OELP (Fig. 1I–I”), whereas Mesh remained in the cytoplasm of the control *hoka*-mutant OELP without Hoka-GFP expression (Fig. 1G–H”). These observations indicate that Hoka is responsible for sSJ organization.

### Hoka is a novel sSJ-protein

To determine the expression pattern and the subcellular localization of endogenous Hoka, we used two anti-Hoka antibodies that were raised against the C-terminal cytoplasmic region of Hoka. In a western blot analysis, the anti-Hoka antibodies detected an intense ~21 kDa band in the extracts from whole wild-type first-instar larvae (Fig. S2A). The ~21 kDa band was absent in *hoka*-mutant extracts (Fig. S2A), indicating that the ~21 kDa band represents Hoka. Immunofluorescence microscopy analyses revealed that an anti-Hoka antibody (29-1) labeled the midgut and the apicolateral region of the OELP in late-stage embryos (Fig. 2A, A’, S2F). The staining pattern of the OELP with the antibody overlapped that of the anti-Mesh antibody (Fig. 2A’, S2F–F”). Furthermore, the immunoreactivity of the antibody in the OELP and midgut was reduced in the *hoka*-mutant embryos (Fig. S2G–I), demonstrating the specificity of the anti-Hoka antibody. Immunofluorescence staining of the first-instar larvae revealed honeycomb-like signals for Hoka in the midgut, OELP, and Malpighian tubules, but not in the foregut and hindgut (Fig. 2B, C). At a higher magnification, staining with the anti-Hoka antibody overlapped with that of the anti-Mesh antibody in the apicolateral region of the midgut epithelial cells (Fig. 2E–E”). The anti-Hoka antibody also labeled the cell-cell contacts in adult midgut epithelial cells (Fig. 2D) and coincided with the staining of the anti-Mesh antibody in the apicolateral region (Fig. 2F–F”). Taken together, these observations indicate that Hoka is a component of sSJs in *Drosophila* from the embryo to adulthood.

**Figure 2.**
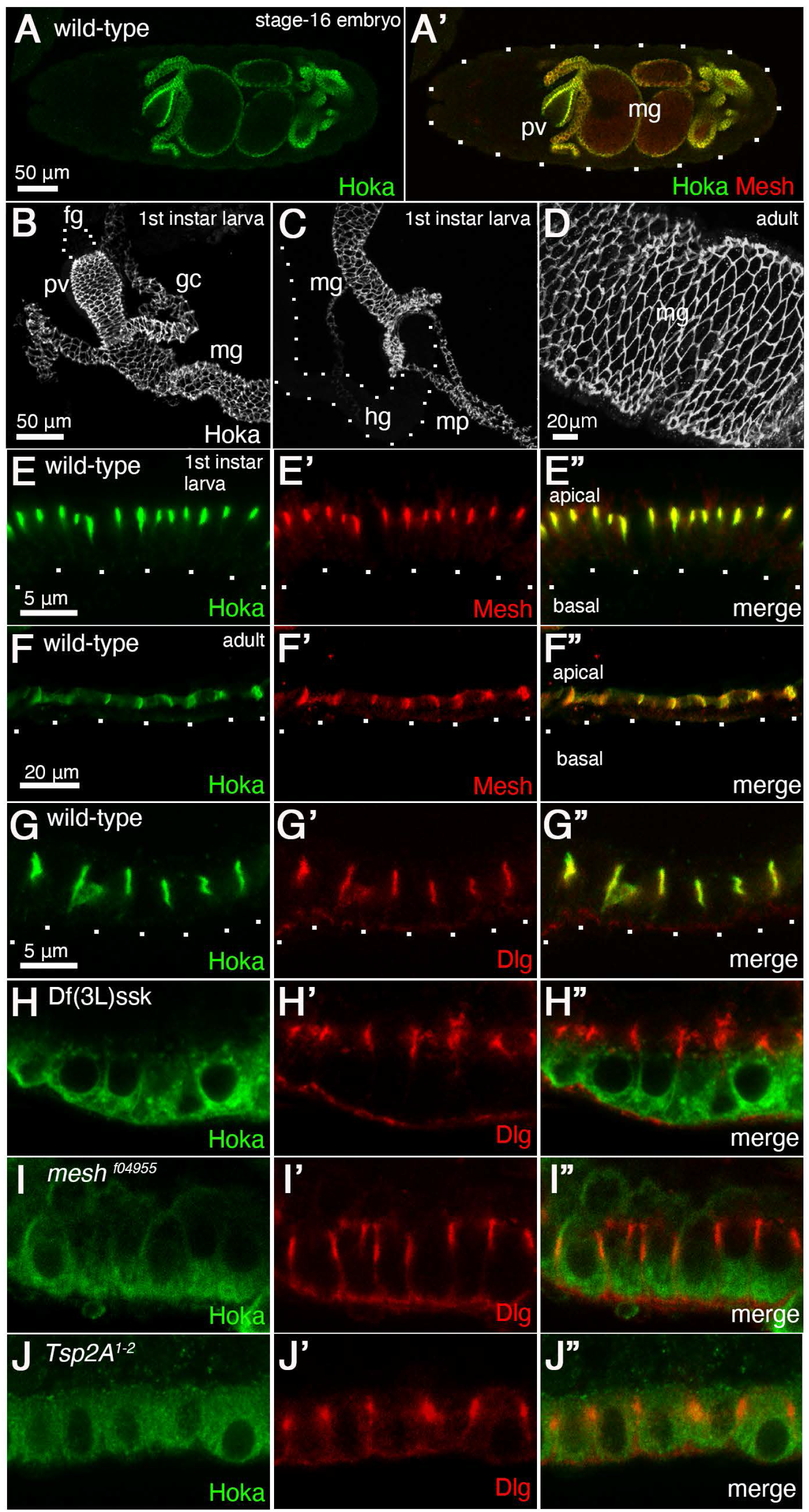
Hoka localizes to sSJs. **(A, A’)** Immunofluorescence staining of a stage 16 wild-type embryo using anti-Hoka (A, A’) and anti-Mesh (A’) antibodies. pv, proventriculus; mg, midgut. The outline of the embryo is delineated by dots. Scale bar: 50 μm. **(B–D)** Immunofluorescence staining of the wild-type first-instar larval anterior midgut (B), posterior midgut (C), and adult midgut (D) using an anti-Hoka antibody. Hoka is expressed in the first-instar larval midgut, the OELP, and the Malpighian tubules (B, C). Hoka signals were not detected in the foregut (B) or hindgut (C). fg, foregut; pv, proventriculus; gc, gastric caeca; mg, midgut; mp, Malpighian tubules; hg, hindgut. Scale bar: 50 μm (B, C), 20 μm (D). **(E–F”)** Immunofluorescence staining of the wild-type first-instar larval (E–E”) and adult (F–F”) midgut using anti-Hoka (green in E, F) and anti-Mesh (red in E’, F’) antibodies. Basal membranes are delineated by dots. Scale bar: 5 μm (E), 20 μm (F). **(G–J”)** The first-instar larval midgut of wild-type, Df(3L)ssk, *mesh^f04955^*-mutant, and *Tsp2A^1-2^*-mutant stained with anti-Hoka (green in G–J) and anti-Dlg (red in G’–J’) antibodies. Scale bar: 5 μm.

We next investigated whether the localization of Hoka is affected by the loss of Mesh, Ssk, or Tsp2A. In the Df(3L)ssk, *mesh^f04955^*, and *Tsp2A^1-2^* mutant first-instar larval midgut epithelial cells, Hoka failed to localize to the apicolateral region but was distributed diffusely and formed aggregates in the cytoplasm (Fig. 2H, I, J), although Dlg was present in the apicolateral region (Fig. 2 H’, I’, J’). Thus, Ssk, Mesh, and Tsp2A are required for the sSJ localization of Hoka.

### Hoka is required for the initial assembly of sSJ-proteins

We next examined Hoka distribution during sSJ formation using immunofluorescence staining of wild-type embryos from stage 14 to stage 16. In the OELP of stage 14 embryos, Hoka was distributed in the cytoplasm and along the lateral membranes (Fig. S3A), and was localized along the lateral membrane with partial accumulation in the apicolateral region in the stage 15 OELP (Fig. S3B). In the stage 16 OELP, Hoka accumulated at the apicolateral region (Fig. S3C), suggesting that it is incorporated into the sSJs during stage 15 to stage 16 of embryonic development. These signals are specific for Hoka, as they were absent in the *hoka*-mutant (Fig. S3D–F). Notably, the sSJ targeting property of Hoka was similar to that of Mesh during sSJ formation of OELP (Fig. S3A’–C”).

To test whether the *hoka*-mutation affects the initial assembly of sSJ-proteins, we monitored the distribution of Ssk, Mesh, and Tsp2A during sSJ maturation in the OELP of wild-type and *hoka*-mutant embryos. In the wild-type OELP, a faint apicolateral distribution of Ssk, Mesh, and Tsp2A was observed at stage 15 (Fig. S3B’, H, N), and they were detectable in the apicolateral region at stage 16 (Fig. S3C’, I, O). By contrast, in the *hoka*-mutant OELP, Ssk, Mesh, and Tsp2A failed to accumulate in the apicolateral region during stage 15 to stage 16 of development (Fig. S3E’–E”, F’–F”, K, L, Q, R). Together, these results indicate that Hoka is required for the initial assembly of sSJ-proteins in the OELP.

### Hoka is required for efficient localization of sSJ-proteins to the apicolateral region

We next observed the distribution of sSJ-proteins in the *hoka*-mutant larval OELP and midgut. As reported previously, Dlg, Ssk, Mesh, and Tsp2A are present in sSJs in the wild-type first-instar larval OELP and midgut (Yanagihashi et al., 2012; Izumi et al., 2012; Izumi et al., 2016) (Fig. 3A’, C–E, I’, K–M). Interestingly, in the *hoka*-mutant first-instar larval OELP, Ssk was distributed in the apical and the apicolateral region (Fig. 3F), and Mesh and Tsp2A were present in the apicolateral region (Fig. 3G, H). Dlg was localized at the apicolateral region (Fig. 3B’). These data are in contrast to the observation that Mesh and Tsp2A were distributed diffusely in the cytoplasm, and Ssk was mislocalized to the apical and the lateral membranes in the *hoka*-mutant stage 16 OELP (Fig. S3F’, L, R). In the *hoka*-mutant first-instar larval midgut epithelial cells, Ssk was mislocalized to the apical and lateral membrane (Fig. 3N), and Mesh and Tsp2A were mislocalized along the lateral membrane (Fig. 3O, P). Further, in these larvae, Ssk, Mesh, and Tsp2A were found to be accumulated in the apicolateral region (Fig. 3Q–S). Dlg was localized in the apicolateral region (Fig. 3J’). Taken together, these results indicate that Hoka is required for the efficient localization of Ssk, Mesh, and Tsp2A to the apicolateral region in epithelial cells.

**Figure 3.**
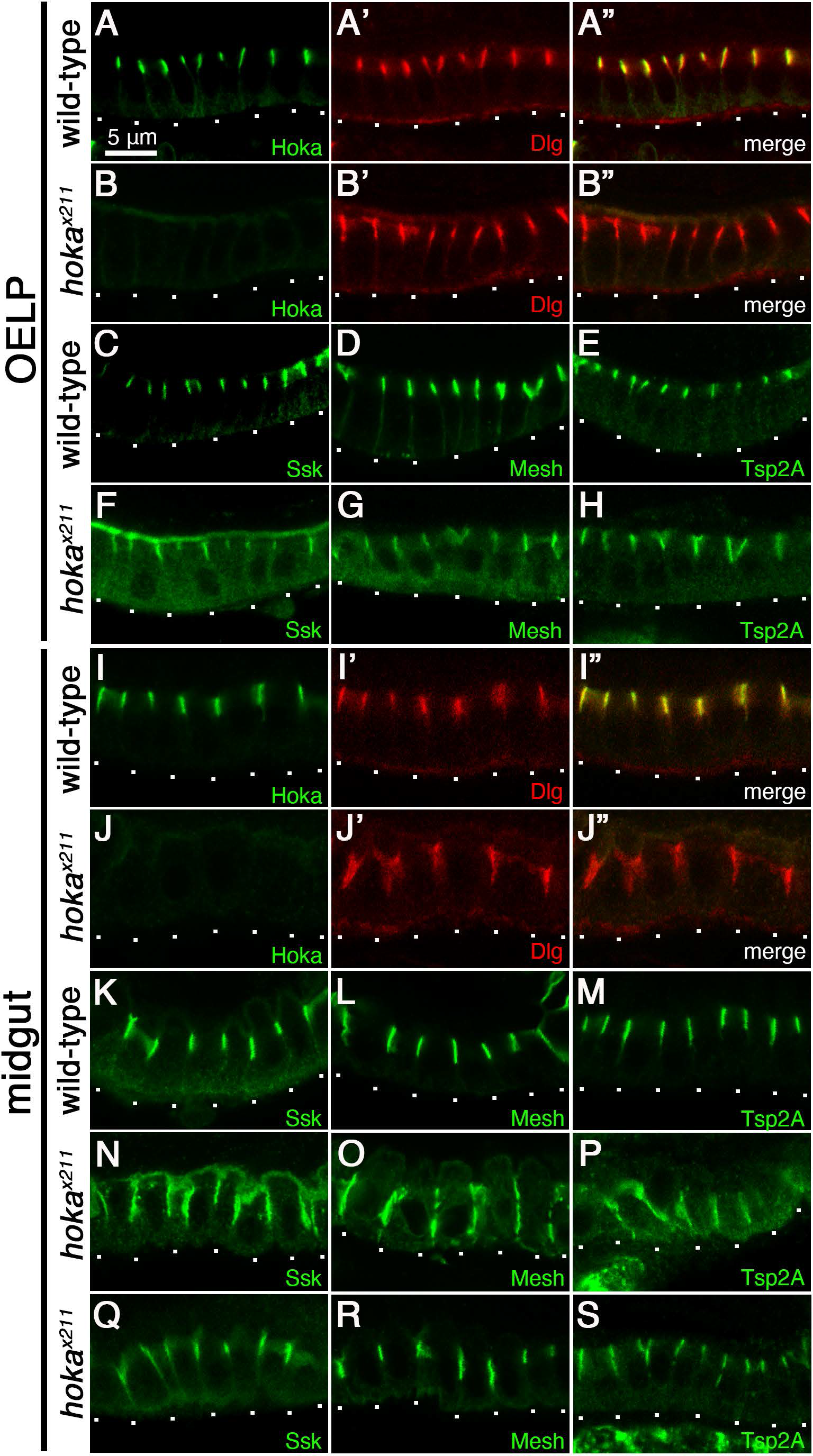
Hoka is required for the localization of sSJ-proteins. **(A–B”)** The first-instar larval OELPs of wild-type (A–A”) and *hoka^x211^*-mutants (B–B”) stained with anti-Hoka (green in A, A”, B, B”) and anti-Dlg (red in A’, A”, B’, B”) antibodies. (**C–H**) The first-instar larval OELPs of wild-type (C, D, E) and *hoka^x211^*-mutant (F, G, H) stained with anti-Ssk (C, F), anti-Mesh (D, G) and anti-Tsp2A (E, H) antibodies. (**I–J”**) The first-instar larval midgut of wild-type (I–I”) and *hoka^x211^*-mutant (J–J”) stained with anti-Hoka (green in I, I”, J, J”) and anti-Dlg (red in I’, I”, J’, J”) antibodies. (**K–S**) The first-instar larval midgut of wild-type (K, L, M) and *hoka^x211^*-mutant (N, O, P, Q, R, S) stained with anti-Ssk (K, N, Q), anti-Mesh (L, O, R) and anti-Tsp2A (M, P, S) antibodies. Basal membranes are delineated by dots. Scale bar (A–S): 5 μm.

Western blot analyses revealed that the densities of the main bands of Ssk (~15 kDa) and Tsp2A (~21 kDa) were not significantly changed in *hoka*-mutant larva, compared with wild-type larvae (Fig. S2B, C). However, the density of Mesh at ~200 kDa and ~90 kDa in the *hoka*-mutant appeared to be increased compared to the wild-type (Fig. S2E), suggesting that Hoka may be involved in the regulation of Mesh protein levels.

### Hoka is required for the proper organization of the sSJ structure

To investigate the role of Hoka in the organization of the sSJ structure, the ultrastructure of the *hoka*-mutant first-instar larval midgut was examined by electron microscopy in ultrathin sections. In the wild-type midgut, sSJs were observed as parallel plasma membranes with ladder-like septa in the apicolateral region of bicellular contacts (Fig. 4A, B). In the *hoka*-mutant midgut, proper sSJ structures were barely detectable at the apicolateral region of bicellular contacts (Fig. 4C–H), although ladder-like structures were occasionally visible (Fig. 4C–E, brackets). Large gaps were often formed between the apicolateral membranes of adjacent cells (Fig. 4F–H, asterisks). Thus, sSJs fail to form correctly in the *hoka*-mutant midgut although Ssk, Mesh, and Tsp2A are present in the lateral regions (Fig. 3N–S). These results indicate that Hoka is required for the proper organization of sSJ structure.

**Figure 4.**
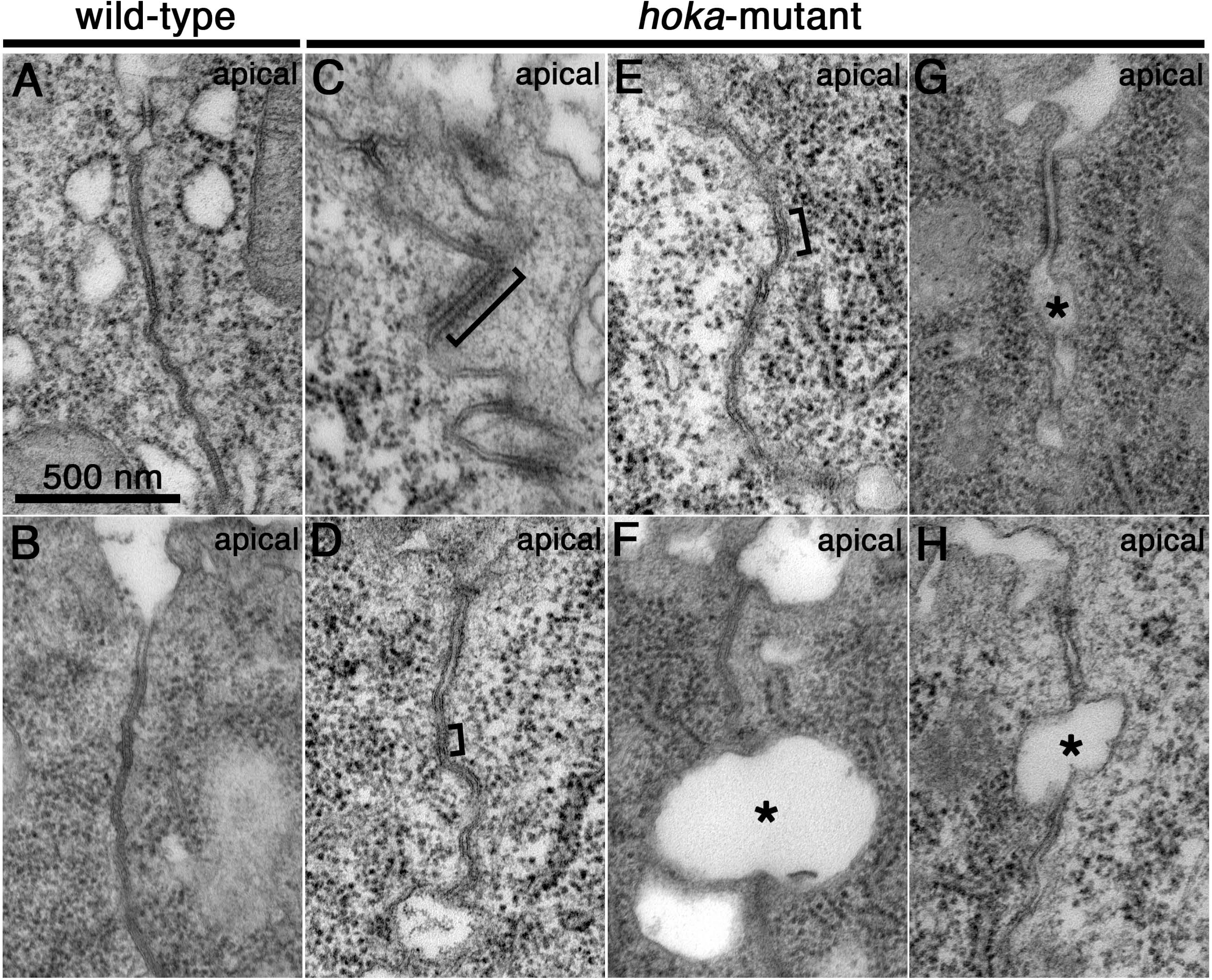
Hoka is required for the correct organization of sSJ structure. **(A–H)** Transmission electron microscopy of the first-instar larval midgut in wild-type (A, B) and *hoka*-mutants (C–H). In the wild-type midgut, typical sSJs are observed at bicellular contacts (A, B). In the *hoka*-mutant midgut, proper sSJ structures are barely detectable at the apicolateral region of bicellular contact (C–H), although the ladder-like structures are occasionally visible (C–E, brackets). Large gaps are often formed between the apicolateral regions of adjacent cells (F–H, asterisks). Scale bar: 500 nm.

### Hoka forms a complex with Ssk, Mesh, and Tsp2A

As Ssk, Mesh and Tsp2A form a complex *in vivo* (Izumi et al., 2012; Izumi et al., 2016), we examined whether Hoka is physically associated with Ssk, Mesh, and Tsp2A. *Drosophila* embryonic extracts were immunoprecipitated using anti-Hoka antibodies, and endogenous Ssk and Mesh were coprecipitated with Hoka (Fig. 5A). Additionally, Hoka was coimmunoprecipitated with Ssk and Mesh from embryonic extracts with anti-Ssk and anti-Mesh antibodies, respectively (Fig. 5B). Neither Hoka, Mesh, nor Ssk was precipitated by the pre-immune sera or the control IgG (Fig. 5A, B). Embryos expressing enhanced GFP (EGFP)-Tsp2A with the *daughterless* (*da*)-GAL4 driver were subjected to immunoprecipitation with anti-GFP antibodies, and EGFP-Tsp2A was found to coprecipitate with endogenous Hoka (Fig. 5C). Hoka was not precipitated from EGFP-Tsp2A-expressing embryos with the control IgG, or EGFP-expressing embryos with the anti-GFP antibody (Fig. 5C). These results indicate that Hoka forms a complex with Ssk, Mesh, and Tsp2A *in vivo*.

**Figure 5.**
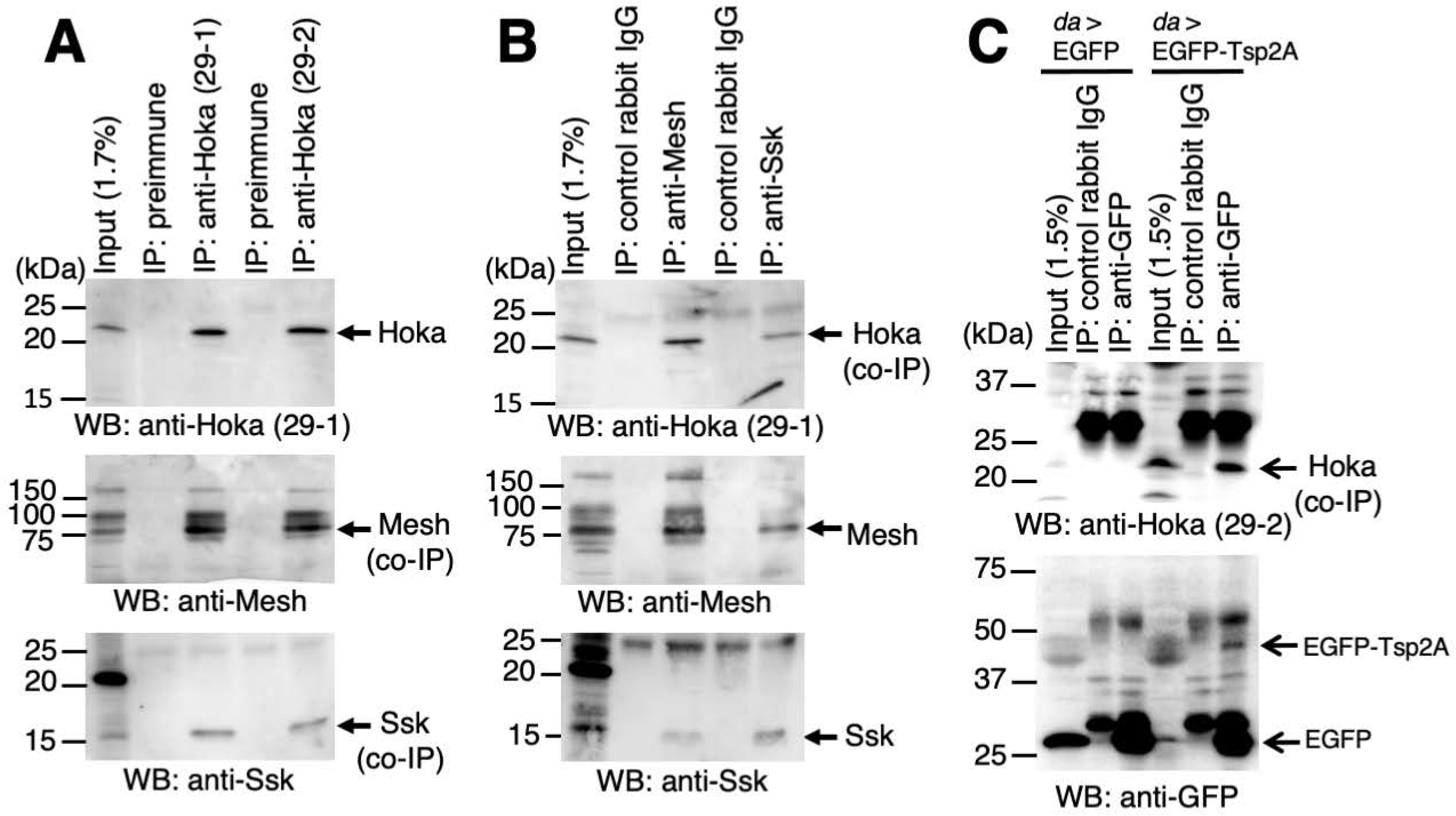
Hoka forms a complex with Ssk, Mesh, and Tsp2A. **(A, B)** Hoka co-immunoprecipitates with Ssk and Mesh. The embryonic extracts (Input) were subjected to immunoprecipitation (IP) with anti-Hoka (A), anti-Mesh (B), and anti-Ssk (B) antibodies. The immunocomplexes were separated on a 15% SDS-polyacrylamide gel, and western blot analyses were performed using anti-Hoka (upper panel), anti-Mesh (middle panel), and anti-Ssk (lower panel) antibodies. Hoka was immunoprecipitated with anti-Hoka antibodies, but not with the pre-immune serum (A, upper panel). The immunoprecipitates of Hoka contained Mesh (A, middle panel) and Ssk (A, lower panel). Mesh was immunoprecipitated with an anti-Mesh antibody, but not with a control IgG (B, middle panel). The immunoprecipitates of Mesh contained Hoka (B, upper panel) and Ssk (B, lower panel). Ssk was immunoprecipitated with an anti-Ssk antibody, but not with a control IgG (B, middle panel). The immunoprecipitates of Ssk contained Hoka (B, upper panel) and Mesh (B, lower panel). **(C)** Hoka co-immunoprecipitates with EGFP-Tsp2A. Extracts of embryos expressing *da*-GAL4/EGFP or *da*-GAL4/EGFP-Tsp2A (Input) were immunoprecipitated (IP) with an anti-GFP antibody. The immunocomplexes were separated on a 15% SDS-polyacrylamide gel, and western blot analyses were performed using anti-Hoka (upper panel) or anti-GFP (lower panel) antibodies. Immunoprecipitations of EGFP-Tsp2A (~50 kDa, arrow) with an anti-GFP antibody are shown (lower panel). EGFP was immunoprecipitated with an anti-GFP antibody from the embryos that expressed EGFP (arrow in lower panel). Hoka was co-precipitated with EGFP-Tsp2A but not with EGFP (arrow in upper panel). Hoka was not precipitated with a control IgG from embryos expressing EGFP-Tsp2A. The kDa indicated on the left-hand side of the images (A–C) refer to the marker band positions.

### Knockdown of *hoka* in the adult ECs leads to increased stem cell proliferation

Recently, we and other groups reported that the knockdown of *ssk*, *mesh*, or *Tsp2A* in adult ECs led to a remarkably shortened lifespan in adult flies, increased ISC proliferation, and intestinal hypertrophy, accompanied by the accumulation of ECs in the midgut (Salazar et al., 2018; Xu et al., 2019; Izumi et al., 2019; Chen et al., 2020). Therefore, we investigated whether the knockdown of *hoka* from adult ECs also caused similar phenotypes as observed with the knockdown of other sSJ-proteins. To knockdown *hoka* in the adult midgut, inducible *hoka*-RNAi was performed using the Gal4/UAS system with an EC-specific driver *Myo1A*-GAL4 and the temperature-sensitive (ts) GAL4 repressor, *tubGal80^ts^* (McGuire et al., 2004; Jiang et al., 2009). *Myo1A-*Gal4 tubGal80^ts^, UAS-*Luciferase* (*Luc*)-RNAi (control), or UAS-*hoka*-RNAi (13704R-1, *hoka*IR-L or *hoka*IR-S; Fig. 1C), flies were raised to adults at 18°C (permissive temperature) and then shifted to 29°C (non-permissive temperature) to inactivate GAL80, leading to the activation of the GAL4/UAS system to express each UAS-driven transgene. Western blot analysis of lysates from the control and the *hoka*-RNAi midgut showed that the Hoka protein level was decreased in the *hoka*-RNAi midgut, compared to the control midgut (Fig. 6A). In the *hoka*-RNAi midgut, Ssk, Mesh, Tsp2A, and Dlg were still observed in the lateral membrane of the ECs (Fig. S4). This is consistent with the observation that Ssk, Mesh, and Tsp2A were distributed to the lateral or apicolateral region in the *hoka*-mutant larval midgut (Fig. 3N–S). Adult flies expressing *hoka*-RNAi had a shortened life span compared to the control flies (Fig. 6B). We also examined whether the barrier function of the midgut was disrupted in *hoka*-RNAi flies. According to the method for a Smurf assay (Rera et al., 2011; Rera et al., 2012), flies were fed a non-absorbable 800-Da blue food dye in sucrose solution. At 5 days after transgene induction, the knockdown of *hoka* in ECs led to a significant increase in flies with blue dye throughout their body cavity, indicating a dysfunction in the midgut barrier (Fig. 6D). The extent of midgut barrier dysfunction in the flies further increased at 8 days after transgene induction, compared with age-matched controls (Fig. 6C, D). Thus, Hoka contributes to the epithelial barrier function in the adult midgut.

**Figure 6.**
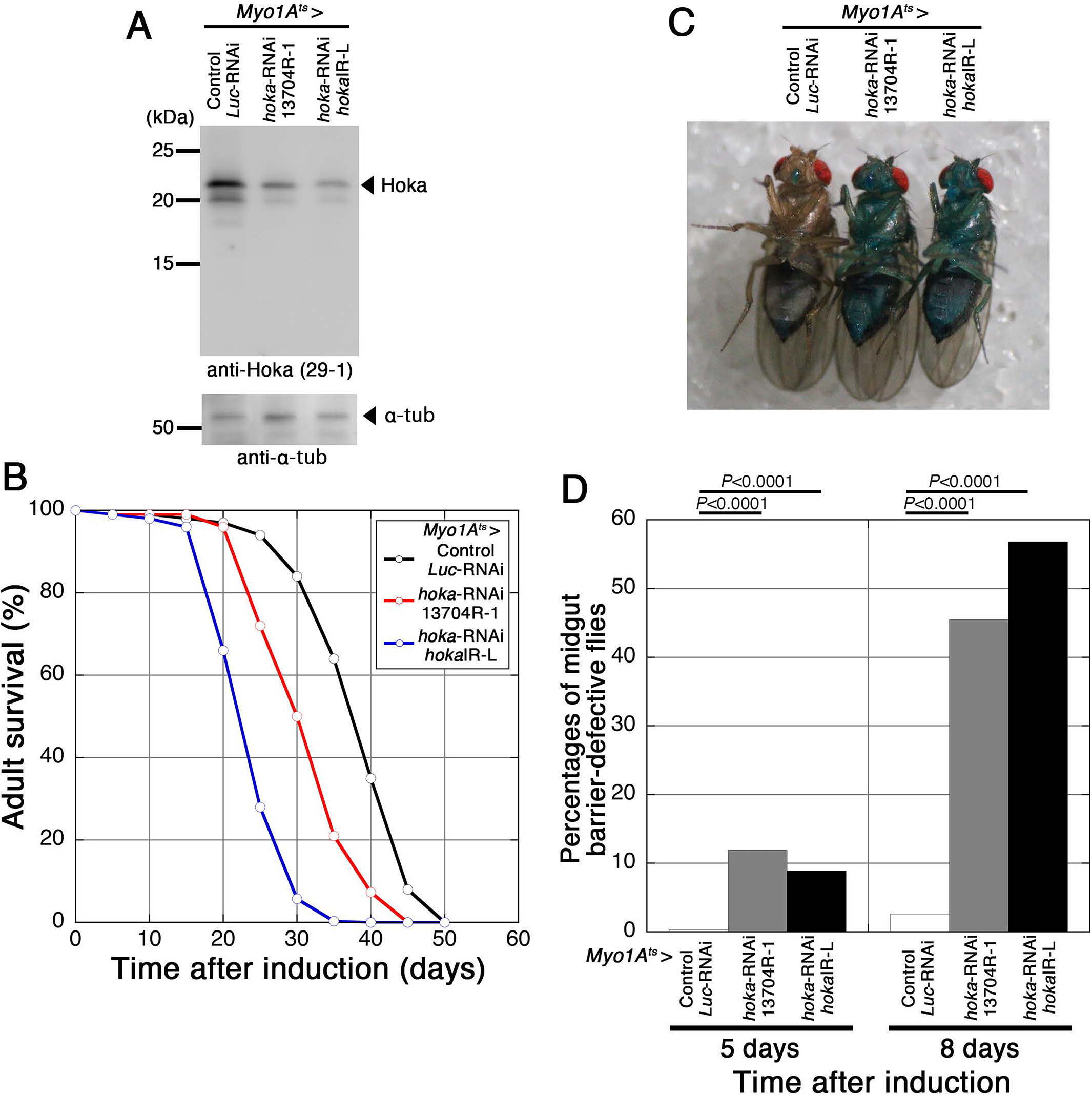
Depletion of *hoka* from ECs in adult flies results in a shortened lifespan and midgut barrier dysfunction. **(A)** Western blot analyses of the *hoka*-RNAi adult midgut. Extracts of the adult midgut prepared from control (*Myo1A^ts^*-Gal4/UAS-*Luc*-RNAi) or *hoka*-RNAi (*Myo1A^ts^*-Gal4/UAS-*hoka*-RNAi 13704R-1 or *hoka*IR-L) flies at 10 days after induction were separated on a 15% SDS-polyacrylamide gel, and western blot analyses were performed using the anti-Hoka (29-1, upper panel) and anti-α-tubulin (lower panel) antibodies. **(B)** Survival analysis of flies expressing *Myo1A^ts^*-Gal4 with UAS-*Luc*-RNAi (control, *n*=300), UAS-*hoka*-RNAi 13704R-1 (*n*=300), or UAS-*hoka*-RNAi *hoka*IR-L (15074R-1) (*n*=300). The transgenes were expressed with GAL80^ts^; therefore, the flies were raised at 18°C until adulthood and were then moved to 29°C. Each vial contained 30 flies (15 females, 15 males). Median lifespan; *Luc*-RNAi: 37 days, *hoka*-RNAi 13704R-1: 30 days, *hoka*IR-L: 22 days. **(C, D)** Barrier integrity (Smurf) assays. Flies expressing *Myo1A^ts^*-Gal4 with UAS-*Luc*-RNAi (control), UAS-*hoka*-RNAi 13704R-1, or UAS-*hoka*-RNAi *hoka*IR-L were fed blue dye in sucrose solution. (C) A control fly and midgut barrier-defective Hoka-deficient flies with blue bodies at 8 days after transgene induction. (D) Left to right: Control (*n*=365), *hoka*-RNAi 13704R-1 (*n*=379), and *hoka*-RNAi *hoka*IR-L (*n*=471) at 5 days after induction, control (*n*=582), *hoka*-RNAi 13704R-1 (*n*=233), and *hoka*-RNAi *hoka*IR-L (*n*=324) at 8 days after induction. The loss of midgut barrier function was determined when the dye was observed outside the midgut. Flies with blue color throughout the body were judged midgut barrier-defective flies although the tone of the color varied depending on the affected flies. *hoka*-RNAi flies showed the loss of barrier function compared with control flies. The *p*-values in (D) represent significant differences in pairwise post-test comparisons indicated by the corresponding bars (Fisher’s exact test).

Next, we examined whether ISC proliferation was increased in the *hoka*-RNAi midgut. Staining the midgut with the phospho-histone H3 (PH3) antibody for the mitotic marker, we found that PH3-positive cells were markedly increased in the *hoka*-RNAi midgut, compared with the midgut in controls (Fig. 7A–C, J). Immunostaining of the midgut with an antibody against Delta, an ISC marker (Ohlstein and Spradling, 2006), showed that ISCs were increased in the *hoka*-RNAi midgut, compared with the midgut in controls (Fig. 7A–C). We also confirmed that the PH3-positive cells were Delta-positive (Fig. 7B, C). The expression of an additional RNAi line for *hoka* (*hoka*IR-S) in ECs also caused increased ISC proliferation (Fig. 7J). These results indicate that knockdown of *hoka* in ECs leads to increased ISC proliferation.

**Figure 7.**
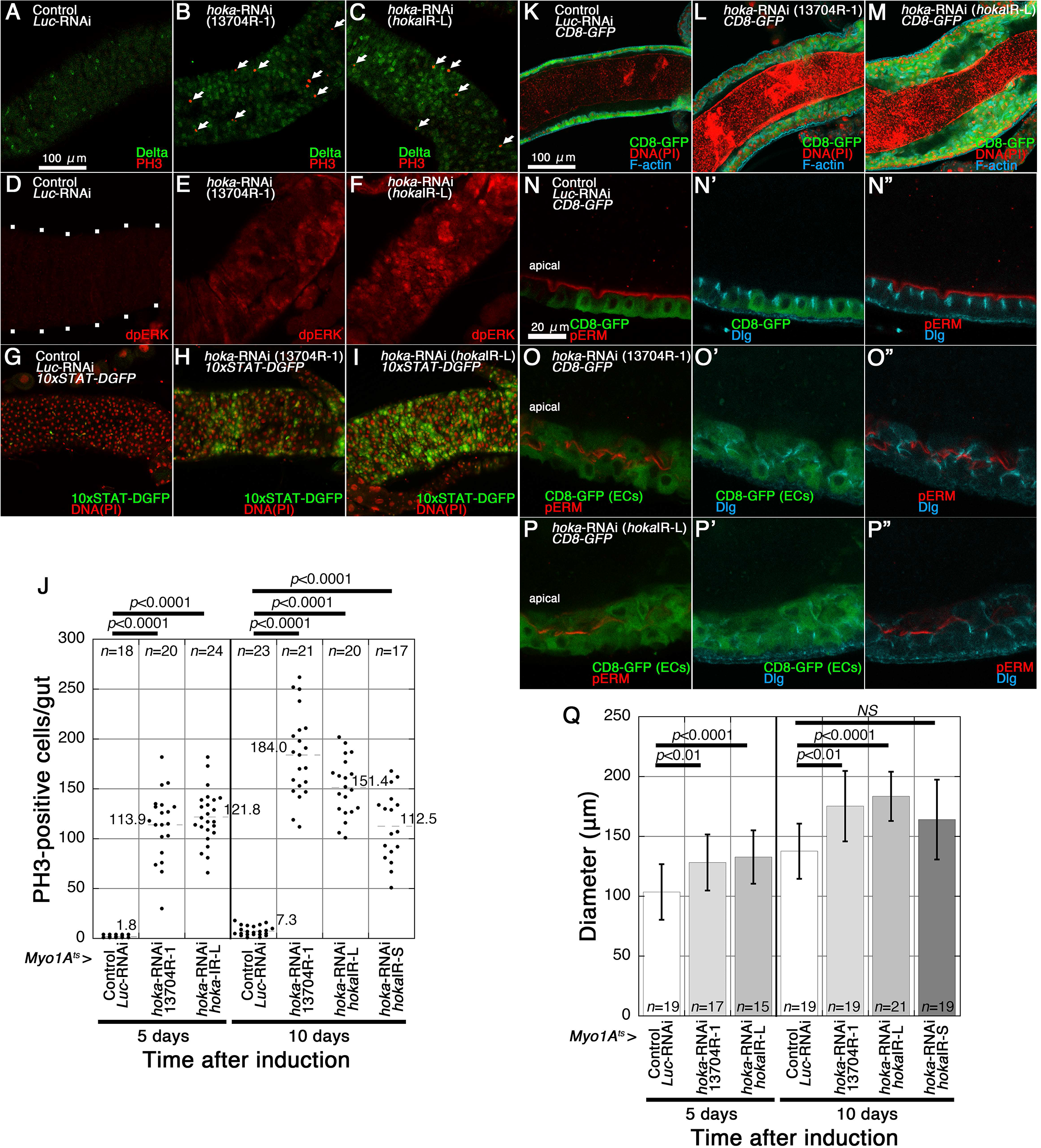
The depletion of *hoka* from ECs leads to increased ISC proliferation and accumulation of ECs in the adult midgut. **(A–C)** Confocal images of the adult posterior midgut expressing *Myo1A^ts^*-Gal4 with UAS-*Luc*-RNAi (control, A), UAS-*hoka*-RNAi 13704R-1 (B), or UAS-*hoka*-RNAi *hoka*IR-L (C) at 10 days after induction and stained for PH3 (red, arrows), and Delta (green). The images show the surface views of the midgut. **(D–F)** Confocal images of the adult posterior midgut expressing *Myo1A^ts^*-Gal4 with UAS-*Luc*-RNAi (control, D), UAS-*hoka*-RNAi 13704R-1 (E), or UAS-*hoka*-RNAi *hoka*IR-L (F) at 10 days after induction and stained for dpERK (red). The images show the surface views of the midgut. The enhancement of Ras-MAPK pathway activity in the *hoka*-RNAi midgut is shown by the increased expression of dpERK (E, F). The outline of the midgut is delineated by dots (D). **(G–I)** Confocal images of the adult posterior midgut expressing *Myo1A^ts^*-Gal4/*10xSTAT-DGFP* with UAS-*Luc*-RNAi (control, G), UAS-*hoka*-RNAi 13704R-1 (H), or UAS-*hoka*-RNAi *hoka*IR-L (I) at 10 days after induction and stained for GFP (green) and DNA (propidium iodide) (red). The enhancement of the Jak-Stat pathway activity in the *hoka*-RNAi midgut is shown by the increased expression of the *10xSTAT-DGFP* reporter (H, I). The images show the surface views of the midgut. Scale bar (A–I): 100 μm. **(J)** Quantification of PH3-positive cells. The dot-plots show the numbers of PH3-positive cells in the individual midguts. Left to right: Control (*n*=18), *hoka*-RNAi 13704R-1 (*n*=20) and *hoka*-RNAi *hoka*IR-L (*n*=24) at 5 days after induction, Control (*n*=23), *hoka*-RNAi 13704R-1 (*n*=21), *hoka*-RNAi *hoka*IR-L (*n*=20), and *hoka*-RNAi *hoka*IR-S (*n*=17) at 10 days after induction. The bars and numbers in the graph represent the mean PH3-positive cells in the fly lines. Statistical significance (*p*<0.0001) was evaluated by one-way ANOVA/Tukey’s multiple comparisons tests. **(K–M)** Confocal images of the adult posterior midgut expressing *Myo1A^ts^*-Gal4/UAS-*CD8-GFP* with UAS-*Luc*-RNAi (control, K), UAS-*hoka*-RNAi 13704R-1 (L), or UAS-*hoka*-RNAi *hoka*IR-L (M) at 10 days after induction and stained for CD8-GFP (green), DNA (propidium iodide) (red), and F-actin (blue). The images show longitudinal cross-sections through the center of the midgut. CD8-GFP driven by *Myo1A^ts^* was expressed in the ECs of each midgut. Scale bar: 100 μm. **(N–P”)** Confocal images of the adult posterior midgut expressing *Myo1A^ts^*-Gal4/UAS-*CD8-GFP* with UAS-*Luc*-RNAi (control, N–N”), UAS-*hoka*-RNAi 13704R-1 (O–O”), or UAS-*hoka*-RNAi *hoka*IR-L (P–P”) at 10 days after induction and stained for pERM (red in N, N”, O, O”, P, P”) and Dlg (blue in N’, N”, O’, O”, P’, P”). The images show the longitudinal cross-sections through the center of the midgut. CD8-GFP driven by *Myo1A^ts^* was expressed in the ECs of each midgut. Scale bar: 20 μm. **(Q)** The diameter of the posterior region of the midgut. The diameter of the midgut was measured just anterior to the Malpighian tubules. Left to right: Control (*n*=19), *hoka*-RNAi 13704R-1 (*n*=17), and *hoka*-RNAi *hoka*IR-L (*n*=15) at 5 days after induction, Control (*n*=19), *hoka*-RNAi 13704R-1 (*n*=19), *hoka*-RNAi *hoka*IR-L (*n*=21), and *hoka*-RNAi *hoka*IR-S (*n*=19) at 10 days after induction. Error bars indicate the SEM. Statistical significance (*p*<0.0001) was evaluated using one-way ANOVA/Tukey’s multiple comparisons tests.

During the adult midgut epithelial regeneration, the Ras-MAP kinase and the Jak-Stat signaling pathways are involved in increased ISC proliferation (Beebe et al., 2010; Buchon et al., 2010; Karpowicz et al., 2010; Shaw et al., 2010; Biteau and Jasper, 2011; Jiang et al., 2009; Jiang et al., 2011; Osman et al., 2012; Zhou et al., 2013). These pathways are activated in the *ssk*, *mesh*, and *Tsp2A*-deficient midgut (Izumi et al., 2019). Therefore, we observed whether these signaling pathways were activated in the *hoka*-RNAi midgut. To monitor the Ras-MAP kinase pathway activity, we examined the levels of diphosphorylated ERK (dpERK) (Gabay et al., 1997). In control flies, dpERK signals were barely detectable in the midgut (Fig. 7D). In contrast, intense dpERK signals were found in the *hoka*-RNAi midgut (Fig. 7E, F), indicating that the Ras-MAP kinase pathway was activated in ISCs. To monitor the Jak-Stat pathway activity, we used a Stat92E reporter line to drive the expression of the destabilized green fluorescent protein (DGFP) (10xSTAT-DGFP). In the control midgut, a few DGFP-positive cells were observed (Fig. 7G), whereas DGFP-positive cells were markedly increased in the *hoka*-RNAi midgut (Fig. 7H, I). Collectively, these results demonstrate that the knockdown of *hoka* in ECs results in the activation of both the Ras-MAP kinase and the Jak-Stat signaling pathways in the midgut.

We next evaluated the organization of the *hoka*-RNAi midgut epithelium. At 10 days after transgene induction, a simple epithelium in which ECs expressed CD8-GFP driven by *Myo1A*-Gal4 was observed in the control midgut (Fig. 7K). The organization of the epithelium was disrupted in the *hoka*-RNAi midgut where several ECs accumulated in the posterior midgut lumen (Fig. 7L, M). In the posterior part of the midgut, the diameter was significantly expanded, compared to the control midgut (Fig. 7Q). The ECs exhibited a variety of aberrant appearances, implying a polarity defect (Fig. 7N–P”). Aberrant distribution of the apical membrane marker phospho-Ezrin/Radixin/Moesin (pERM) (Chen et al., 2018) and Dlg, a marker for the apicolateral membrane, was observed in the *hoka*-RNAi ECs (Fig. 7O–P”). Thus, knockdown of *hoka* in ECs causes intestinal tumor accompanied by the accumulation of ECs in the midgut lumen, indicating that Hoka is required for maintaining intestinal homeostasis in the adult fly. We performed *hoka*-RNAi in ISCs/EBs using an *escargot^ts^* (*esg^ts^*)-GAL4 driver and observed increased ISC proliferation and accumulation of ECs in the midgut (Fig. S5), suggesting that Hoka function is required for ISC and/or EBs to regulate ISC behavior in the adult midgut.

### aPKC and Yki are involved in ISC overproliferation caused by *hoka*-RNAi

In a recent study, the reduced expression of aPKC and the Hippo transcriptional coactivator Yki in *Tsp2A*-RNAi ISCs/EBs or ECs led to the reduction of *Tsp2A*-RNAi-induced ISC overproliferation in the midgut (Xu et al., 2019). Xu et al. also showed that the expression of *Tsp2A*-RNAi in the midgut increases aPKC staining in the cell border membrane. We examined whether aPKC staining was increased in the *Myo1A^ts^*-GAL4-driven *hoka*-RNAi midgut. In the control midgut, aPKC staining was barely detectable, but the signal intensity of aPKC staining was significantly increased in the *hoka*-RNAi midgut, compared to the control midgut (Fig. 8A–C). In the longitudinal cross-sections of the control midgut, apical membrane aPKC staining was occasionally observed in cells with a small nucleus (presumably ISCs) (Fig. 8D–D,” G–G,” arrowhead). aPKC has been reported to localize asymmetrically in the apical membrane of ISCs and regulates the differentiation of ISCs to EBs (Goulas et al., 2012). Interestingly, in the *hoka*-RNAi midgut, apical membrane staining of aPKC was often found in the cells mounted by other cells (Fig. 8E–E,” F–F,” H–H,” arrow). There are large and small nuclei-containing cells in the apical aPKC-localizing cells (Fig. 8E–E”, F–F”, H–H”). The apical aPKC staining partially overlapped with F-actin staining (Fig. 8E”, F”) and Dlg staining (Fig. 8H”). To clarify the cell types in which apical aPKC was observed, we expressed CD8-GFP together with *hoka*-RNAi using the *Myo1A^ts^-GAL4* in the midgut and stained the midgut with an anti-Delta antibody. Here, GFP and Delta-positive cells were identified as ECs and ISCs, respectively. In the control midgut, apical aPKC staining was found in the Delta-positive ISCs (Fig. 8I–I”). In the *hoka*-RNAi midgut, three types of apical aPKC-localizing cells were observed (Fig. 8J–J”, K–K”): the Delta-positive ISCs (arrowheads), the Delta and CD8-GFP-negative cells (presumably EBs and/or EEs) (arrows), and the Delta-negative and CD8-GFP-positive cells (EC-like cells) (yellow arrows). These results indicate that aPKC can be apically localized in ISCs and the differentiated cells in the *hoka*-RNAi midgut.

**Figure 8.**
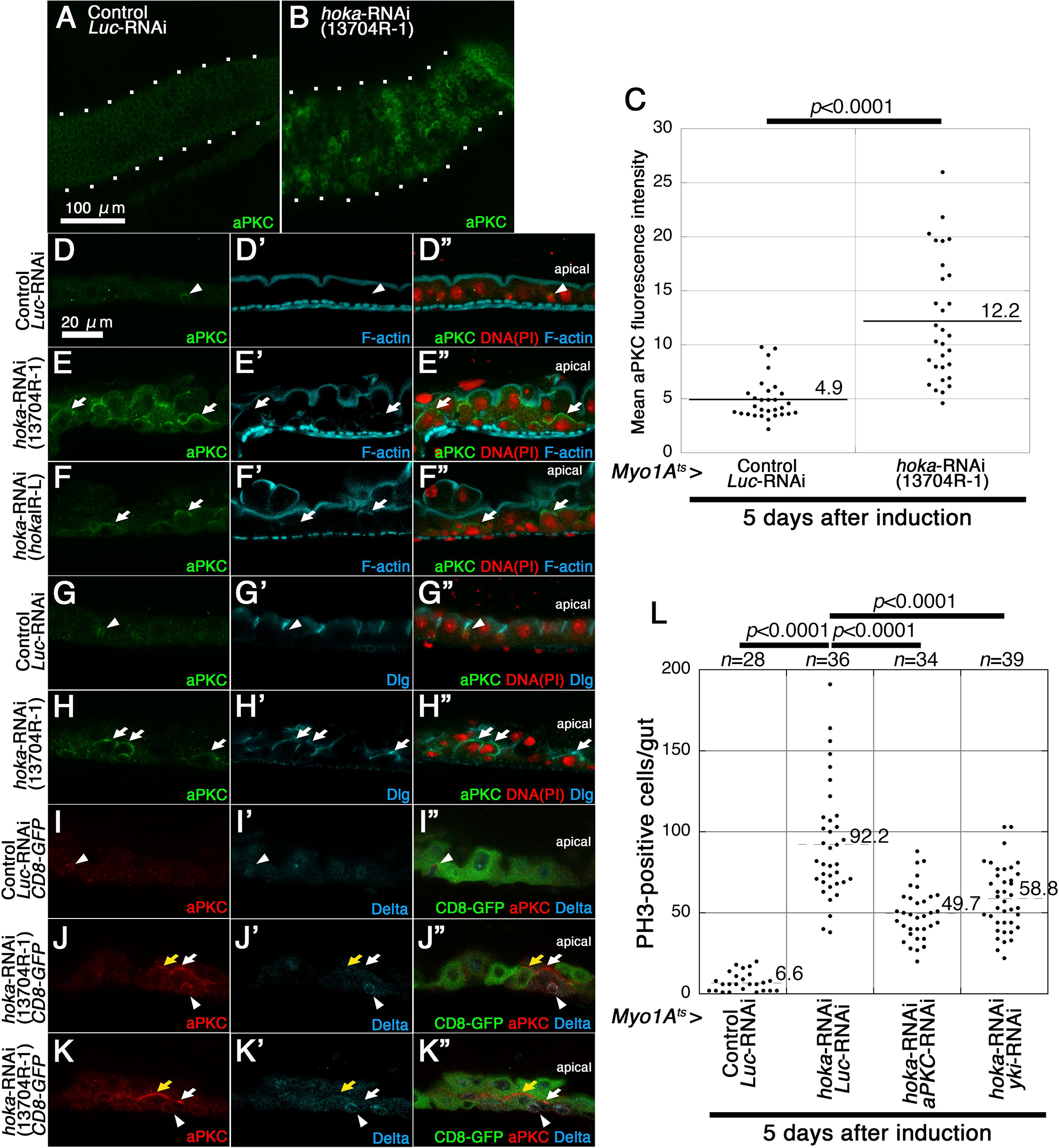
The depletion of *aPKC* and *yki* from *hoka*-RNAi ECs results in the reduction of ISC overproliferation caused by *hoka*-RNAi. **(A, B)** Confocal images of the adult posterior midgut expressing *Myo1A^ts^*-Gal4 with UAS-*Luc*-RNAi (control, A) or UAS-*hoka*-RNAi 13704R-1 (B) at 5 days after induction and stained for aPKC (green). The images show the surface views of the midgut. The outline of the midgut is delineated by dots. Scale bar: 100 μm. **(C)** Dot-plots showing the mean aPKC fluorescence intensity in the posterior midgut. The bars and the numbers in the graph display the mean fluorescence intensity of the control (*Luc*-RNAi) or *hoka*-RNAi 13704R-1 midgut. The mean fluorescence intensity was calculated from three random (100 μm × 100 μm) fields per midgut (*n*=10 for each genotype). Statistical significance (*P*<0.001) was determined using the Mann-Whitney *U* test. **(D–H”)** Confocal images of the adult posterior midgut expressing *Myo1A^ts^-Gal4* with UAS-*Luc*-RNAi (control, D–D”, G–G”), UAS-*hoka*-RNAi 13704R-1 (E–E”, H–H”), or UAS-*hoka*-RNAi *hoka*IR-L (F–F”) at 5 days after induction and stained for aPKC (green in D–H, D”–H”), F-actin (blue in D’–F’, D”–F”), and Dlg (blue in G’, G”, H’, H”). The arrowheads indicate the cells with apical aPKC staining and a small nucleus in the control midgut. The arrows indicate cells with apical aPKC staining and a large nucleus in the *hoka*-RNAi midgut. The images show the longitudinal cross-sections through the center of the midgut. **(I–K”)** Confocal images of the adult posterior midgut expressing *Myo1A^ts^*-Gal4/UAS-*CD8-GFP* with UAS-*Luc*-RNAi (control, I–I”) or UAS-*hoka*-RNAi 13704R-1 (J–K”) at 5 days after induction and stained for aPKC (red in I, I”, J, J”, K, K”) and Delta (blue in I’, I”, J’, J”, K’, K”). The arrowheads indicate Delta and aPKC double-positive cells. The arrows and the yellow arrows indicate the apical aPKC-positive and CD8-GFP-negative cells, and the apical aPKC and CD8-GFP-double positive cells, respectively. CD8-GFP driven by *Myo1A^ts^* was expressed in the ECs of each midgut. The images show the longitudinal cross-sections through the center of the midgut. Scale bar (D–K”): 20 μm. **(L)** Quantification of PH3-positive cells. The dot-plots show the numbers of PH3-positive cells in the individual midguts. Left to right: Control (*n*=28), *hoka*-RNAi 13704R-1 together with *Luc*-RNAi (*n*=36), *hoka*-RNAi 13704R-1 together with *aPKC*-RNAi HMS01411 (*n*=34), and *hoka*-RNAi 13704R-1 together with *yki*-RNAi JF03119 (*n*=39) at 5 days after induction. Bars and numbers in the graph represent the mean PH3-positive cells in the fly lines. Statistical significance (*p*<0.0001) was evaluated by one-way ANOVA/Tukey’s multiple comparisons tests.

Next, we investigated whether the depletion of *aPKC* and *yki* from *hoka*-RNAi ECs results in a reduction in ISC overproliferation caused by *hoka*-RNAi. To deplete the expression of *aPKC* and *yki* in *hoka*-RNAi ECs, we used *aPKC* (HMS01411) and *yki* (JF03119) RNAi lines, both of which efficiently reduced ISC overproliferation caused by the *Tsp2A*-RNAi (Xu et al., 2019). Expression of *hoka*-RNAi together with *aPKC*-RNAi or *yki*-RNAi by *Myo1A^ts^*-GAL4 in ECs significantly reduced ISC overproliferation, compared to the *hoka*-RNAi and the *Luc*-RNAi midgut (Fig. 8L, S6A–D). Accumulation of cells in the midgut lumen was still observed in the *hoka*-RNAi together with *aPKC-RNAi* or *yki*-RNAi midguts (Fig. S6E–H), probably due to the high rate of ISC proliferation in these midguts (Fig. 8L). Taken together, these results indicate that aPKC and Yki activities mediate ISC overproliferation caused by *hoka*-RNAi in the ECs.

## Discussion

The identification of Ssk, Mesh, and Tsp2A has provided an experimental system to analyze the role of sSJs in the *Drosophila* midgut (Furuse and Izumi, 2017). Recent studies have shown that sSJs regulate the epithelial barrier function and also ISC proliferation and EC behavior in the midgut (Salazar et al., 2018; Xu et al., 2019; Izumi et al., 2019; Chen et al., 2020). Furthermore, sSJs are involved in epithelial morphogenesis, fluid transport, and macromolecule permeability in the Malpighian tubules (Jonusaite et al., 2020; Beyenbach et al., 2020). Here, we have reported the identification of a novel sSJ-associated membrane protein Hoka. Hoka is required for the efficient accumulation of other sSJ-proteins at sSJs and the correct organization of sSJ structure. The knockdown of *hoka* in the adult midgut leads to intestinal barrier dysfunction, increased ISC proliferation mediated by aPKC and Yki activities, and epithelial tumors. Thus, Hoka contributes to sSJ organization and maintaining ISC homeostasis in the *Drosophila* midgut.

### The unique primary structure of Hoka homolog proteins

Arthropod sSJs have been classified together based on their morphological similarity (Green and Bergquist, 1982; Lane, 1994). The identification of sSJ proteins in *Drosophila* has provided an opportunity to investigate whether sSJs in various arthropod species share similarities at the molecular level. However, Hoka homolog proteins appear to be conserved only in insects upon a database search (data not shown), suggesting compositional variations in arthropod sSJs.

Interestingly, the cytoplasmic region of Hoka includes three YTPA motifs. The same or similar amino acid motifs are also present in the Hoka homologs of other insects, such as other *Drosophila* species, mosquitos, and a butterfly (YQPA motif) although the number of these motif(s) vary (1 to 3 in *Drosophila* species, 1 in mosquitos, 1 in a butterfly). The extensive conservation of the YTPA/YQPA motif in insects suggests that the motif plays a critical role in the molecular function of Hoka homologs. It would be interesting to investigate the role of the YTPA/YQPA motif in sSJ organization.

The extracellular region of Hoka appears to be composed of 13 amino acids alone after the cleavage of the signal peptide, which is too short to bridge the 15–20 nm intercellular space of sSJs (Lane, 1994; Tepass and Hartenstein, 1994). Thus, Hoka is unlikely to act as a cell adhesion molecule in sSJs. Indeed, the overexpression of Hoka-GFP in *Drosophila* S2 cells did not induce cell aggregation, which is a criterion for cell adhesion activity (data not shown).

### The role of Hoka in sSJ organization

The loss of an sSJ-protein results in the mislocalization of other sSJ-proteins (Izumi et al., 2012; Izumi et al., 2016), indicating that sSJ-proteins are mutually dependent for their sSJ localization. In the *ssk*-deficient midgut, Mesh and Tsp2A were distributed diffusely in the cytoplasm (Izumi et al., 2012; Izumi et al., 2016). In the *mesh*-mutant midgut, Ssk was localized at the apical and lateral membranes, whereas Tsp2A was distributed diffusely in the cytoplasm (Izumi et al., 2012; Izumi et al., 2016). In the *Tsp2A*-mutant midgut, Ssk was localized at the apical and lateral membranes, whereas Mesh was distributed diffusely in the cytoplasm (Izumi et al., 2016). Among these three mutants, the mislocalization of Ssk, Mesh, or Tsp2A is consistent; Mesh and Tsp2A were distributed in the cytoplasm, whereas Ssk was localized at the apical and lateral membranes. However, in the *hoka*-mutant larval midgut, Mesh and Tsp2A were distributed along the lateral membrane, whereas Ssk was mislocalized to the apical and lateral membranes. Interestingly, in some *hoka*-mutant midguts, Ssk, Mesh, and Tsp2A were localized to the apicolateral region, as observed in the wild-type midgut. Differences in subcellular misdistribution of sSJ-proteins between the *hoka*-mutant and the *ssk*, mesh, and *Tsp2A*-mutants indicate that the role of Hoka in the process of sSJ formation is different from that of Ssk, Mesh, or Tsp2A. Ssk, Mesh, and Tsp2A may form the core complex of sSJs, and these proteins are indispensable for the generation of sSJs, whereas Hoka facilitates the arrangement of the primordial sSJs at the correct position, i.e., the apicolateral region. This Hoka function may also be important for rapid paracellular barrier repair during the epithelial cell turnover in the adult midgut. Notably, during the sSJ formation process of OELP, the sSJ targeting property of Hoka was similar to that of Mesh, implying that Hoka may have a close relationship with Mesh, rather than Ssk and Tsp2A during sSJ development.

### The role of Hoka in intestinal homeostasis

The knockdown of *hoka* in the adult midgut leads to a shortened lifespan in adult flies, intestinal barrier dysfunction, increased ISC proliferation, and the accumulation of ECs. These results are consistent with the recent observation for *ssk*, *mesh*, and *Tsp2A*-RNAi in the adult midgut (Salazar et al., 2018; Xu et al., 2019; Izumi et al., 2019; Chen et al., 2020). However, the defects observed in the *hoka*-RNAi midgut were less severe than in flies with RNAi for other sSJ-proteins (Izumi et al., 2019). At 5 days after RNAi induction, only ~10% of *hoka*-RNAi adult flies showed a midgut barrier defect, whereas more than 45% of flies with RNAi for other sSJ-proteins exhibited the barrier defect (Izumi et al., 2019). The median lifespan of *hoka*-RNAi adult flies (13704R-1: 30 days, *hoka*IR-L: 22 days) was much longer than in flies with RNAi for other sSJ-proteins (*ssk*-RNAi: 7 days, *mesh*-RNAi, 12074R-1: 7 days, *Tsp2A*-RNAi, 11415R-2; 8 days) (Izumi et al., 2019). Additionally, EC accumulation in *hoka*-RNAi flies was modest compared to flies with RNAi for other sSJ-proteins, where a large number of ECs fill the posterior midgut lumen 5 days after RNAi induction (Izumi et al., 2019). These modest defects in the *hoka-*RNAi midgut may reflect an insufficient reduction of Hoka expression. However, given that other sSJ-proteins are still present at the lateral membrane in the *hoka*-mutant larval and *hoka*-RNAi adult midgut, and the septa-like structures were often observed in the bicellular contacts in the *hoka*-mutant larval midgut, sSJ function may be partly maintained in the *hoka*-deficient midgut.

The intestinal barrier dysfunction caused by RNAi for sSJ-proteins may permit the leakage of particular substances from the midgut lumen, which may induce particular cells to secrete cytokines and growth factors for ISC proliferation. Alternatively, sSJs or sSJ-associated proteins may be directly involved in the secretion of cytokines and growth factors through the regulation of intracellular signaling in the ECs. In the latter case, Xu et al. (2019) showed that *Tsp2A* knockdown in ISCs/EBs or ECs hampers the endocytic degradation of aPKC, thereby activating the aPKC and Yki signaling pathways, leading to ISC overproliferation in the midgut. Therefore, Xu et al. proposed that sSJs are directly involved in the regulation of aPKC and the Hippo pathway-mediated intracellular signaling for ISC proliferation. We have shown that the expression of *hoka*-RNAi together with *aPKC*-RNAi or *yki*-RNAi in ECs significantly reduced ISC overproliferation caused by *hoka*-RNAi. Thus, aPKC- and Yki-mediated ISC overproliferation appears to commonly occur in sSJ-protein-deficient midguts. However, the possibility that the leakage of particular substances through the paracellular route may be involved in ISC overproliferation in the sSJ-proteins-deficient midgut cannot be excluded.

It has been reported that apical aPKC staining is observed in ISCs but is barely detectable in ECs (Goulas et al., 2012). We found that the expression of *hoka*-RNAi in ECs increased aPKC staining in the midgut. Additionally, in the *hoka*-RNAi midgut, apical aPKC staining was observed in ISCs and in differentiated cells including EC-like cells. Thus, apical and increased cytoplasmic aPKC may contribute to ISC overproliferation. Interestingly, EC-like cells in the *hoka*-RNAi midgut do not always localize aPKC to the apical regions. Apical aPKC staining was detected in EC-like cells mounted by other cells but was barely detectable in the lumen-facing EC-like cells. These mounted cells are thought to be newly generated cells after the induction of *hoka*-RNAi, which may not be able to exclude aPKC from the apical region in the crowded cellular environment. A recent study showed that aberrant sSJ formation caused by *Tsp2A*-depletion impairs aPKC endocytosis and increases aPKC localization in the membrane of cell borders (Xu et al., 2019). The sSJ-proteins including Hoka may also regulate endocytosis to exclude aPKC from the apical membrane of ECs.

aPKC is a key determinant of apical-basal polarity in various epithelia (Ohno et al., 2015). However, in the *Drosophila* adult midgut, aPKC is not required for the establishment of the epithelial cell polarity (Chen et al., 2018). Instead, its activity contributes to ISC overproliferation caused by the depletion of sSJ-proteins in the midgut. Therefore, it is intriguing that the roles of aPKC in the adult midgut are different from those in several ectoderm derived epithelia. The identification of molecules involved in aPKC-mediated ISC proliferation may provide a better understanding of the aPKC-mediated signaling pathway as well as the mechanisms underlying the increased expression and apical targeting of aPKC in the ECs deficient for sSJ-proteins. It would be of particular interest for future studies to analyze whether cell-cell junction dysfunction triggers aPKC activation to regulate stem cell proliferation in metazoan tissues other than the *Drosophila* adult midgut.

## Materials and methods

### Fly stocks and genetics

Fly stocks were reared on a standard cornmeal fly medium at 25°C. *w^1118^* flies were used as wild-type flies unless otherwise specified. The other fly stocks used were *w*, *da*-GAL4/TM6B, *Tb* (#55851; Bloomington *Drosophila* Stock Center (BDSC), Bloomington, IN), *y w*; *Myo1A*-GAL4 (#112001; *Drosophila* Genetic Resource Center (DGRC), Kyoto, Japan), *tubP*-GAL80^*ts*^ (#7019; BDSC), *y w; Pin^Yt^*/CyO; UAS-*mCD8*-GFP (#5130; BDSC), *w*; *10xStat92E-DGFP*/TM6C *Sb Tb* (#26200; BDSC), *y w*; *esg-lacZ*/CyO (#108851; DGRC), and FRT19A; *ry* (#106464; DGRC). The RNAi lines used were *hoka*-RNAi (#13704R-1, Fly Stocks of National Institute of Genetics (NIG-Fly), Mishima, Japan), *Luciferase* (*Luc*)-RNAi (#31603; BDSC), *aPKC*-RNAi (HMS01411, #35001; BDSC), *yki*-RNAi (JF03119, #31965; BDSC). The mutant stocks used were Df(3L)ssk (Yanagihashi et al., 2012), *mesh^f04955^* (#18826; BDSC) (Izumi et al., 2012), *Tsp2A^1-2^* (Izumi et al., 2016). For the phenotype rescue experiment, pUAST vectors (Brand and Perrimon, 1993) containing *hoka-GFP* were constructed and a fly strain carrying this construct was established. The stocks used for the generation of *hoka*-mutants were *y^2^ cho^2^ v^1^* (TBX-0004), *y^1^ v^1^* P{*nos-phiC31/int.NLS*}X; attP40 (II) (TBX-0002; NIG-Fly), *y^2^ cho^2^ v^1^*; Sco/CyO (TBX-0007; NIG-Fly), *y^2^ cho^2^ v^1^* P{*nosP-Cas9, y^+^, v^+^*}1A/FM7c, *Kr*GAL4 UAS-GFP (CAS0002; NIG-Fly), and *y^2^ cho^2^ v^1^; PrDr*/TM6C, *Sb Tb* (TBX-0010; NIG-Fly).

### Antibodies

The following antibodies were used: rabbit-anti-Hoka (29-1; 1:2000, 29-2; 1:2000), rabbit anti-Mesh (955-1; 1:1000), rabbit anti-Mesh (995-1; 1:1000), rat anti-Mesh (8002; 1:500) (Izumi et al., 2012), rabbit anti-Ssk (6981-1; 1:1000) (Yanagihashi et al., 2012), rabbit anti-Tsp2A (302AP, 1:200) (Izumi et al., 2016), mouse anti-Dlg (4F3, Developmental Studies Hybridoma Bank (DSHB); 1:50), mouse anti-Delta (C594.9B; DSHB; 1:20), rabbit anti-PH3 (06-570; Millipore, Darmstadt, Germany; 1:1000), rabbit anti-dpERK (4370; Cell Signaling, Danvers, MA; 1:500), rabbit anti-phospho-Ezrin/Radixin/Moesin (pERM) (3726; Cell Signaling, 1:200), rat anti-GFP (GF090R; Nakalai Tesque; 1:1000), rabbit anti-GFP (598; MBL, Nagoya, Japan; 1:1000), rabbit anti-aPKC (sc-216; Santa Cruz, Dallas, TX; 1:500). Alexa Fluor 488-conjugated (A21206; Invitrogen), and Cy3- and Cy5-conjugated (712-165-153 and 715-175-151, Jackson ImmunoResearch Laboratories, West Grove, PA, USA) secondary antibodies were used at 1:400. Actin was stained with Alexa Fluor 568 phalloidin (A12380; Thermo Fisher; 1:1000) or Alexa Fluor 647 phalloidin (A22287; Thermo Fisher; 1:1000). Nuclei were stained with propidium iodide (Nakalai Tesque; 0.1 mg ml^-1^).

### Deficiency screen

Embryos of deficiency lines were obtained from the *Drosophila* Stock Center and the *Drosophila* Genetic Resource Center (Kyoto, Japan). The deficiency screen was performed as described previously (Izumi et al., 2016).

### cDNA cloning and expression vector construction

The ORF of *hoka*, including the initiation codon, was amplified by PCR with the forward primer harboring an EcoRI site (5’-cggaattcACGAAAACGACGGAAATGAAGTTGGCT-3’) and the reverse primer having a BglII site (5’-gaagatctGTGACAATAGCGGTGGCATGCG-3’), and the EcoRI and BglII sites containing *Drosophila* embryonic cDNA was cloned into the EcoRI and BglII sites of the pUAST vector (Brand and Perrimon, 1993). To generate an expression vector for the C-terminal EGFP-tagged Hoka, EGFP cDNA with 3’ and 5’ XhoI sites was cloned into the XhoI site of pUAST-*hoka*. To generate RNAi lines (*hoka*IR-L and *hoka*IR-S), a DNA fragment containing 1 to 682 (*hoka*IR-L) or 160 to 682 (*hoka*IR-S) of the *hoka* ORF was amplified by PCR with the forward primer harboring an EcoRI site (*hoka*IR-L; 5’-cggaattcATGAAGTTGGCTAAGAAGTGC-3’, *hoka*IR-S; 5’-cggaattcATCGTTTGTGTAGCGGTAGGT-3’) or a XhoI site (*hoka*IR-L; 5’-ccgctcgagATGAAGTTGGCTAAGAAGTG-3’, *hoka*IR-S; 5’-ccgctcgag ATCGTTTGTGTAGCGGTAGGT-3’), and the reverse primer having a BglI site (5’-gaagatctTCAGACAATAGCGGTGGCATG-3’). The two types of DNA fragments were inserted into pUAST as a head-to-head dimer and transformed into SURE2 competent cells (200152; Agilent Technologies, Santa Clara, CA, USA). Transgenic flies were generated by standard P-element transformation procedures.

### Generation of *Tsp2A* mutants

Generation of *hoka* mutants using the CRISPR/Cas9 system was performed according to the method described by Kondo and Ueda (Kondo and Ueda, 2013). To construct a guide RNA (gRNA) expression vector for *hoka*, two complementary 24 bp oligonucleotides of the target sequence with 4 bp overhangs on both ends (5’-cttcGGCCTGCTGCCTGCAAGAAT-3’ and 5’-aaacATTCTTGCAGGCAGCAGGCC-3’) were annealed to generate a double-stranded DNA fragment which was cloned into BbsI-digested pBFv-U6.2 (NIG-Fly) (pBFv-U6.2-*hoka*CR1). pBFv-U6.2-*hoka*CR1 was injected into the *y^1^ v^1^* nos-phiC31; attP40 host (Bischof et al., 2007). Surviving G_0_ males were individually crossed to *y^2^ cho^2^ v^1^* virgins. A single male transformant from each cross was mated to *y^2^ cho^2^ v^1^*; Sco/CyO virgins. Offspring in which the transgene was balanced were collected to establish a stock. Males carrying a U6.2-*hoka*CR1 transgene were crossed to *nos*-Cas9 females (*y^2^ cho^2^ v^1^* P {*nosP-Cas9, y^+^, v^+^*}1A) to obtain founder flies that have both the U6.2-*hoka*CR1 and the *nos*-Cas9 transgenes. Male founders were crossed to *y^2^ cho^2^ v^1^; PrDr*/TM6C, *Sb Tb* female flies. Each male possessing *y^2^ cho^2^ v^1^*/Y; +/ TM6C, *Sb Tb* genotype was crossed to *y^2^ cho^2^ v^1^*; *PrDr*/TM6C, *Sb Tb* female flies and the offspring possessing the *y^2^ cho^2^ v^1^*; +/ TM6C, *Sb Tb* genotype were collected to establish the lines. The embryos of the lethal lines were immunostained for Hoka. Genomic DNA of the *hoka*-negative lines was extracted and analyzed for mutations in the *hoka* gene locus.

### Production of polyclonal antibodies against Hoka

The amino acids 77-136 encoding the cytoplasmic region of the Hoka protein were cloned into pGEX-6P (GE Healthcare) to produce a GST (Glutathione S-transferase) fusion protein. The proteins were expressed in *Escherichia coli* (DH5α). Polyclonal antibodies were generated in rabbits (29-1 and 29-2 by Kiwa Laboratory Animals (Wakayama, Japan)).

### Immunostaining

Embryos were fixed with 3.7% formaldehyde in PBS for 20 min. Adult flies and larvae were dissected in Hanks’ Balanced Salt Solution and the midgut was fixed with 4% paraformaldehyde in PBS/0.2% Tween-20 for 30 min. The fixed specimens were washed thrice with PBS/0.4% Triton X-100 and were blocked with 5% skim milk in PBS/0.2% Tween-20. Thereafter, the samples were incubated with primary antibodies at 4°C overnight, washed thrice with PBS/0.2% Tween-20, and incubated with secondary antibodies for 3 h. After another three washes, the samples were mounted in Fluoro-KEEPER antifade reagent (12593-64; Nakalai Tesque, Kyoto, Japan). Images were acquired with a confocal microscope (Model TCS SPE; Leica Microsystems, Wetzlar, Germany) using its accompanying software and the HC PLAN Apochromat 20× NA 0.7 and HCX PL objective lenses (Leica Microsystems). Images were processed with Adobe Photoshop® software (Adobe Systems Inc., San Jose, CA).

### Electron microscopy

First-instar larvae of wild-type or *hoka^x211^*-mutants were dissected and fixed overnight at 4°C with a mixture of 2.5% glutaraldehyde and 2% paraformaldehyde in 0.1 M cacodylate buffer (pH 7.4). The specimens including the midguts were prepared as described previously (Izumi et al., 2012). Ultrathin sections (50–100 nm) were stained doubly with 4% hafnium (IV) chloride and lead citrate and observed with a JEM-1010 electron microscope (JEOL, Tokyo, Japan) equipped with a Veleta TEM CCD Camera (Olympus, Tokyo, Japan) at an accelerating voltage of 80 kV.

### Co-immunoprecipitation and western blotting

Fly wild-type embryos and embryos expressing EGFP-Tsp2A or GFP (*da*-Gal4>UAS-EGFP-Tsp2A or UAS-GFP) were mixed with a 5-fold volume of lysis buffer (30 mM Tris-HCl pH 7.5, 150 mM NaCl, 1% Brij97 (P6136, Sigma-Aldrich, St. Louis, MO, USA)) and protease inhibitor cocktail (25955-11, Nakarai Tesque, Kyoto, Japan) and homogenized using a pestle for 1.5 ml microfuge tubes. The method for immunoprecipitation was essentially the same as described previously (Izumi et al., 2016). Rabbit anti-Hoka (29-1, 29-2), rabbit anti-Mesh (995-1), rabbit anti-Ssk (6981-1), and rabbit anti-GFP (598, MBL, Nagoya, Japan) antibodies were used for immunoprecipitation. Immunocomplexes and extracts of the first-instar larva were separated on SDS-polyacrylamide gels, transferred to polyvinylidene difluoride membranes and western blot analyses were performed using rabbit anti-Hoka (29-1; 1:2000, 29-2; 1:2000), rabbit anti-Mesh (995-1; 1:1000), rabbit anti-Ssk (6981-1; 1:1000), rabbit anti-GFP (598; 1:1000, MBL, Nagoya, Japan), and mouse anti-α-tubulin (DM-1A; 1:1000, Sigma-Aldrich) antibodies. The molecular weights of protein bands were estimated using the Precision Plus Protein™ Dual Color Standards (#161-0374; BIO-RAD, Hercules, CA).

### Conditional expression of UAS transgenes (TARGET system)

Flies were crossed and grown at 18°C until eclosion. Adult female flies were collected 2–5 days after eclosion and transferred to 29°C for the inactivation of Gal80.

### Barrier integrity (Smurf) assay

Flies at 2–5 days of age were placed in empty vials containing a piece of paper soaked in 2.5% (wt/vol) Blue Dye No. 1 (Tokyo Chemical Industry, Tokyo, Japan)/5% sucrose solution at 50–60 flies/vial. After 2 days at 18°C, the flies were placed in new vials containing paper soaked in BlueDye/sucrose and transferred to 29°C. Loss of midgut barrier function was determined when the dye was observed outside the gut (Rera et al., 2011) (Rera et al., 2012). Flies were transferred to new vials every 2 days.

### Quantification of aPKC intensity

To measure the fluorescence intensity of aPKC, a z-projection was created from the R5 region of the posterior midgut. The projection included stacks of the EC layer of the midgut. The mean gray value of the green channel was collected from three random (100 μm × 100 μm) fields per midgut using the ImageJ software 1.52k (U. S. National Institutes of Health, Bethesda, MD, USA) and we subtracted the background values measured from the outside area surrounding the midgut.

### Statistical analyses

Statistical significance was evaluated using the Mann–Whitney *U*-test, Student’s *t*-test, one-way ANOVA/Tukey’s multiple comparisons test (KaleidaGraph software; Synergy Software, Reading, PA), and the Fisher’s exact test. Values of *p*<0.05 were considered significant.

## Abbreviations

aPKC: atypical protein kinase C
DGFP: destabilized green fluorescent protein
Dlg: Discs large
*da*-GAL4: *daughterless*-GAL4
dpERK: diphosphorylated ERK
EB: enteroblast
EC: enterocyte
EGFP: enhanced green fluorescent protein
EE: enteroendocrine cell
EMC: enteroendocrine mother cell
*esg^ts^*-GAL4: *escargot^ts^*-GAL4
ISC: intestinal stem cell
Luc: Luciferase
OELP: outer epithelial layer of the proventriculus
PH3: phospho-histone H3
pERM: phospho-Ezrin/Radixin/Moesin
pSJ: pleated septate junction
SJ: septate junction
sSJ: smooth septate junction
sSJ-proteins: specific molecular constituents of sSJs
Ssk: Snakeskin
ts: temperature-sensitive
Upd: Unpaired
Yki: Yorki

## Acknowledgments

We are grateful to all members of the Furuse laboratory for helpful discussions. We thank the Bloomington Drosophila Stock Center, *Drosophila* Genetic Resource Center at Kyoto Institute of Technology, and Fly Stocks of National Institute of Genetics (NIG-Fly) for fly stocks. We would like to thank Editage (www.editage.com) for their assistance with English language editing.

## Author contributions

Y.I. designed the research; Y.I. and K.F. performed the experiments; Y.I. analyzed the data; Y.I. and M.F. wrote the paper.

---

**Figure S1.**
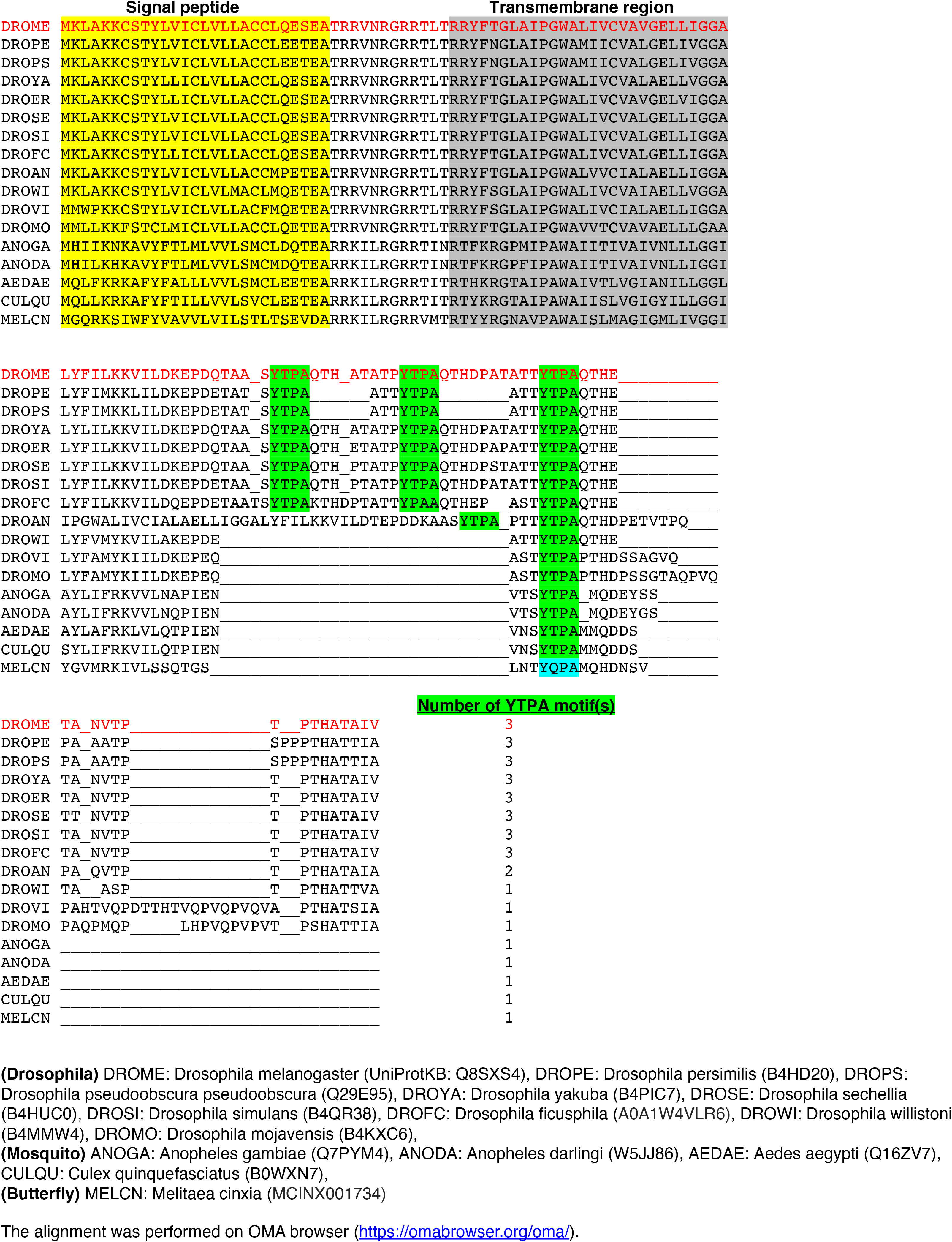
Multiple sequence alignment of the Hoka amino acid sequence with the homologs in *Drosophila*, mosquito, and butterfly. The MAFFT multiple sequence alignment was performed using the OMA browser (https://corona.omabrowser.org/oma/home/). The amino acid sequence of *Drosophila melanogaster* (DEOME) Hoka is shown at the top (red). Signal peptides and transmembrane regions are highlighted in yellow and gray, respectively. YTPA (Tyr-Thr-Pro-Ala) motifs are highlighted by green. Although the Hoka homolog of the butterfly *Melitaea cinxia* (MELCN) does not possess the YTPA motif, a similar motif (YQPA) is found in the cytoplasmic region. The number of YTPA/YQPA motif(s) is indicated at the end of each sequence.

**Figure S2.**
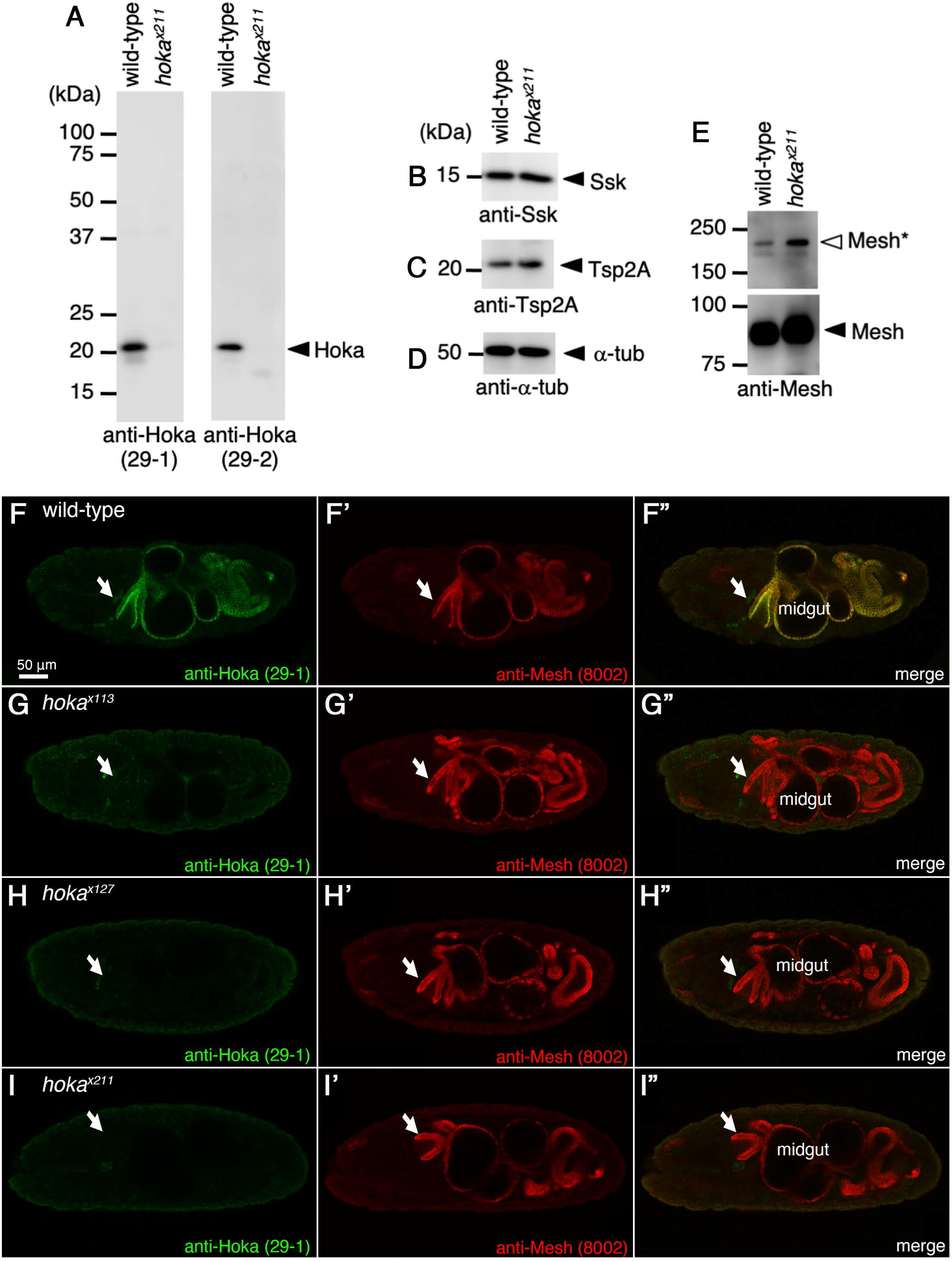
Characterization of the *hoka*-mutant strains and anti-Hoka antibodies. **(A–E)** The extracts from the first-instar larva prepared from wild-type and *hoka^x211^*-mutants were separated on 15% (A–D) or 8% (E) SDS-polyacrylamide gels, and western blot analyses were performed using the anti-Hoka (A, left panel, 29-1; right panel, 29-2), anti-Ssk (B), anti-Tsp2A (C), anti-α-tubulin (D), and anti-Mesh (E) antibodies. A protein band of ~21 kDa was detected by anti-Hoka antibodies in the wild-type but not in the *hoka^x211^-mutant* (E; arrowheads), indicating that the ~21 kDa band represents Hoka. The density of the main bands of Ssk (~15 kDa) and Tsp2A (~21 kDa) were not significantly different in the *hoka^x211^*-mutant relative to the wild-type (F, G; arrowhead). The density of the main band for Mesh was slightly increased in the *hoka^x211^*-mutant compared to the wild-type (E; arrowhead). In the higher-molecular-mass band for Mesh, which showed a double band at ~200 kDa, the upper band (Mesh*) was increased in the *hoka^x211^-mutant* compared to the wild-type (E; white arrowhead). Western blots using the anti-α-tubulin antibody as the loading control show that the same quantities of protein were loaded in the wild-type and *hoka^x211^*-mutant extracts (D). **(F–I”)** Immunofluorescence staining of stage 16 wild-type (F–F”), *hoka^x113^* (G–G”), *hoka^x127^* (H–H”), or *hoka^x211^* (I–I”) embryos using the anti-Hoka (29-1, green) and anti-Mesh (8002, red) antibodies. The immunoreactivity of the anti-Hoka antibody (29-1) was diminished in the *hoka^x113^*, *hoka^x127^*, or *hoka^x211^* embryos. The arrows indicate OELPs. Bars: 50 μm.

**Figure S3.**
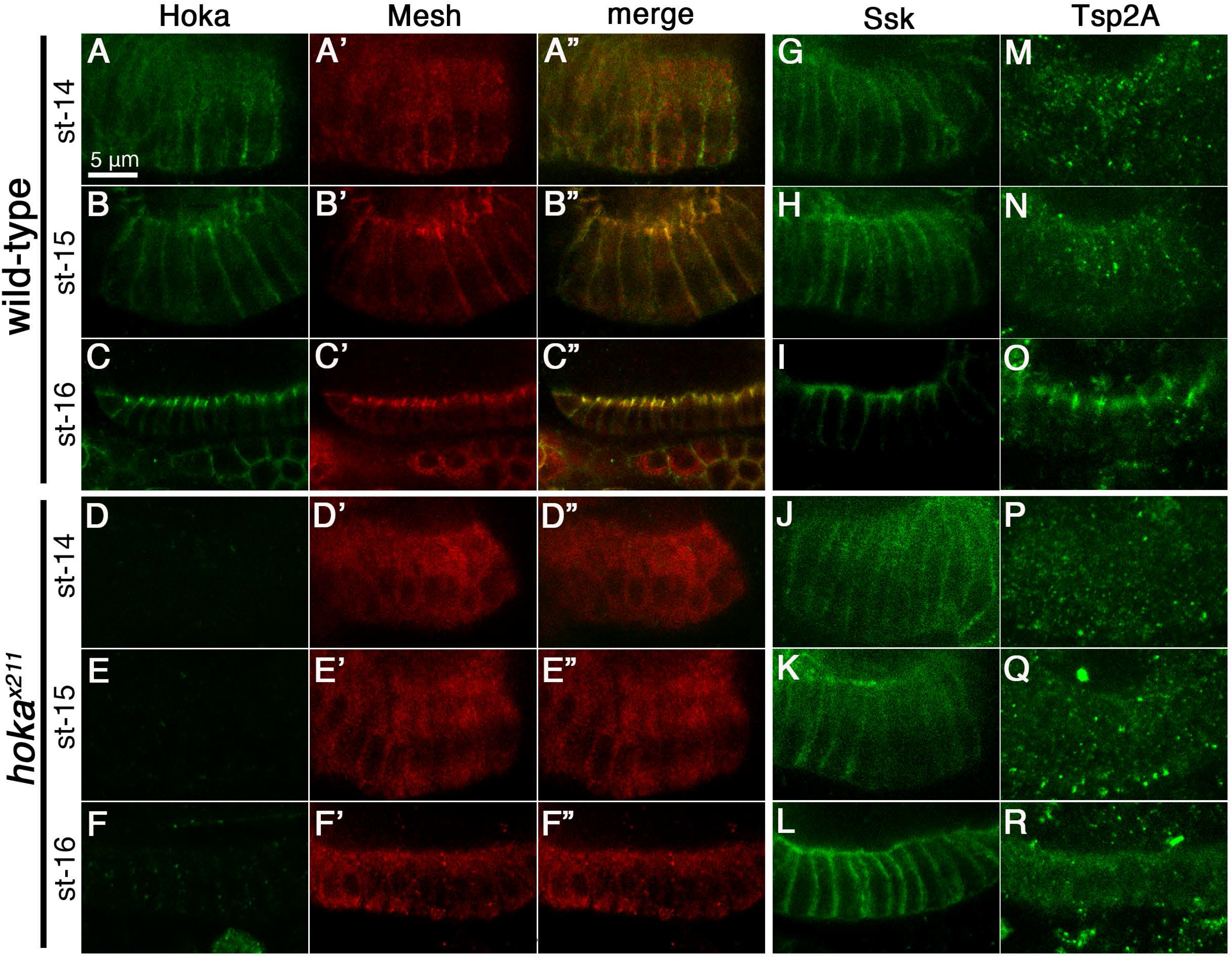
Hoka, Ssk, Mesh, and Tsp2A distribution during sSJ formation in the wild-type and *hoka^x211^*-mutant embryonic OELPs. **(A–R)** Immunofluorescence staining of stage 14 wild-type OELPs (A–A”, G, M), *hoka^x211^*-mutant OELPs (D–D”, J, P), stage 15 wild-type OELPs (B–B” and H, N), *hoka^x211^*-mutant OELPs (E–E”, K, Q), stage 16 wild-type OELPs (C–C”, I, O), and *hoka^x211^*-mutant OELPs (F–F”, L, R) with the anti-Hoka (29-1 for A–F, A”–F”), anti-Mesh (8002 for A’–F’, A”–F”), anti-Ssk (6981-1 for G–L), and anti-Tsp2A (301AP for M–R). Scale bar (A–R): 5 μm.

**Figure S4.**
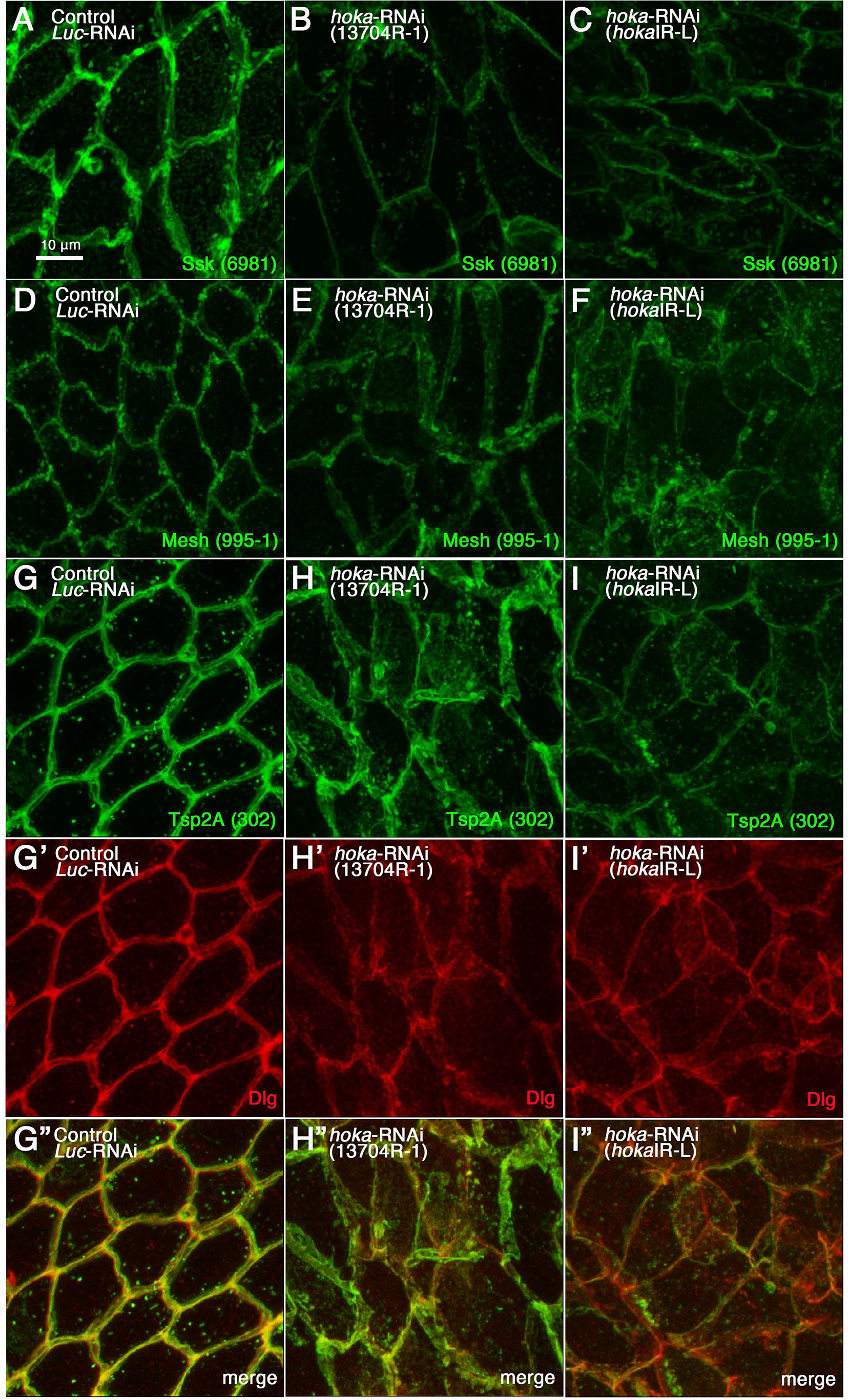
The distribution of sSJ-proteins in the *hoka*-RNAi adult midgut. **(A–I”)** Confocal images of the adult posterior midgut expressing *Myo1A^ts^*-Gal4 with UAS-*Luc*-RNAi (control, A, D, G–G”), UAS-*hoka*-RNAi 13704R-1 (B, E, H–H”), or UAS-*hoka*-RNAi *hoka*IR-L (C, F, I–I”) at 10 days after induction and stained for Ssk (6981-1 for A–C), Mesh (995-1 for D–F), Tsp2A (302 for G–I, G”–I”), Dlg (G’–I’, G”–I”). The images show the surface views of the midgut. Scale bar (A–I”): 10 μm.

**Figure S5.**
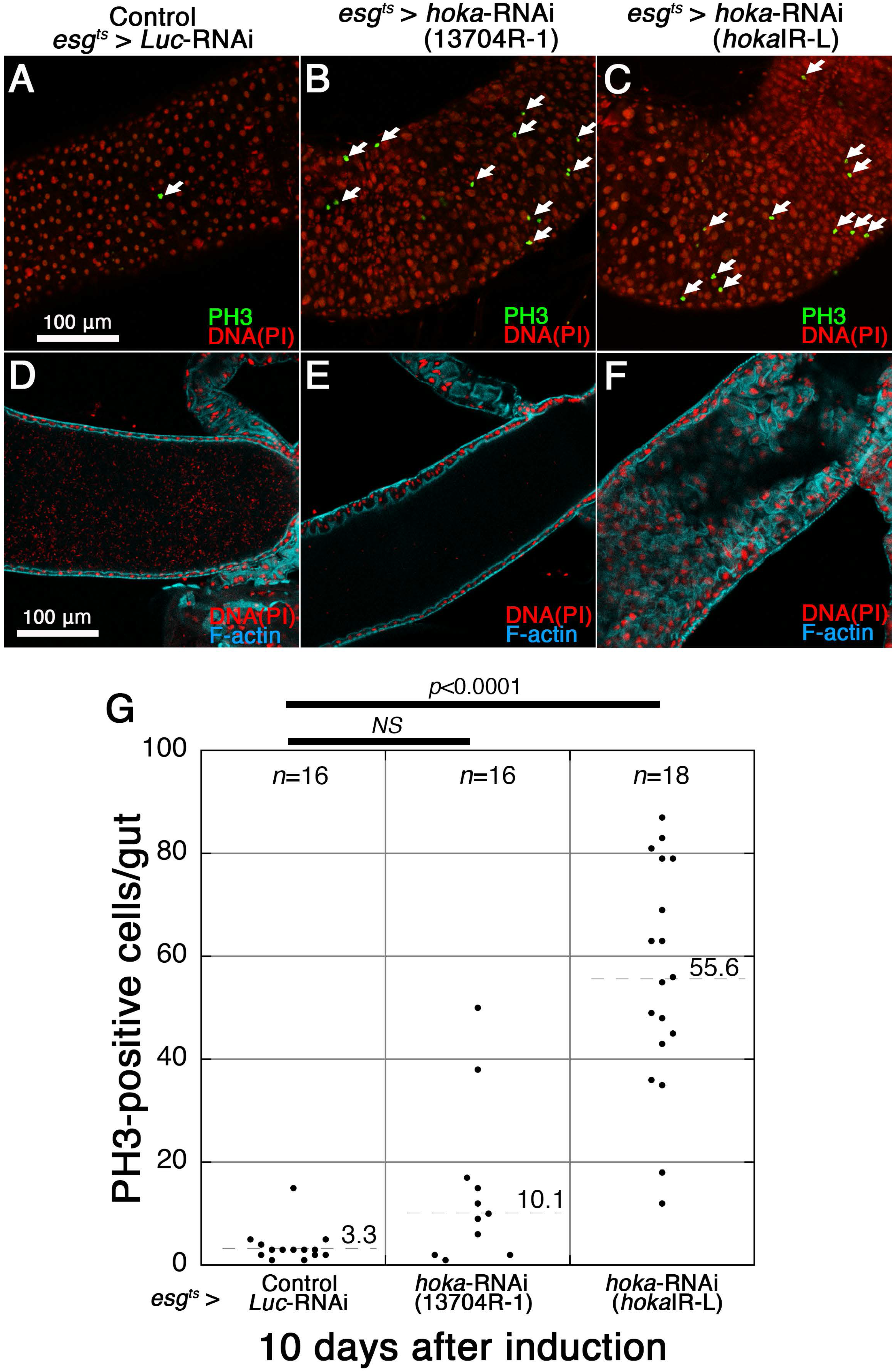
The knockdown of *hoka* in progenitor cells leads to increased ISC proliferation and accumulation of ECs in the adult midgut. **(A–C)** Confocal images of the adult posterior midgut expressing *esg^ts^*-Gal4 with UAS-*Luc*-RNAi (control, A), UAS-*hoka*-RNAi 13704R-1 (B), or UAS-*hoka*-RNAi *hoka*IR-L (C) at 10 days after induction stained for PH3 (green, arrows) and DNA (propidium iodide) (red). The images show the surface views of the midgut. PH3-positive cells were increased in the *hoka*-RNAi midgut compared with the control midgut. Scale bar: 100 μm. **(D–F)** Confocal images of the adult posterior midgut expressing *esg^ts^*-Gal4 with UAS-*Luc*-RNAi (control, D), UAS-*hoka*-RNAi 13704R-1 (E), or UAS-*hoka*-RNAi *hoka*IR-L (F) at 10 days after induction and stained for DNA (propidium iodide) (red) and F-actin (blue). The images show the longitudinal cross-sections through the center of the midgut. Several ECs were accumulated in the *hoka*-RNAi *hoka*IR-L midgut lumen (F). Scale bar: 100 μm. **(G)** Quantification of PH3-positive cells. The dot-plots show the numbers of PH3-positive cells in individual midguts. Left to right: Control (*Luc*-RNA*i*) (*n*=16), *hoka*-RNAi 13704R-1 (*n*=16) and *hoka*-RNAi *hoka*IR-L (*n*=18) at 10 days after induction. The bars and numbers in the graph represent the mean PH3-positive cells in the fly lines. Statistical significance (*p*<0.0001) was evaluated by one-way ANOVA/Tukey’s multiple comparisons tests.

**Figure S6.**
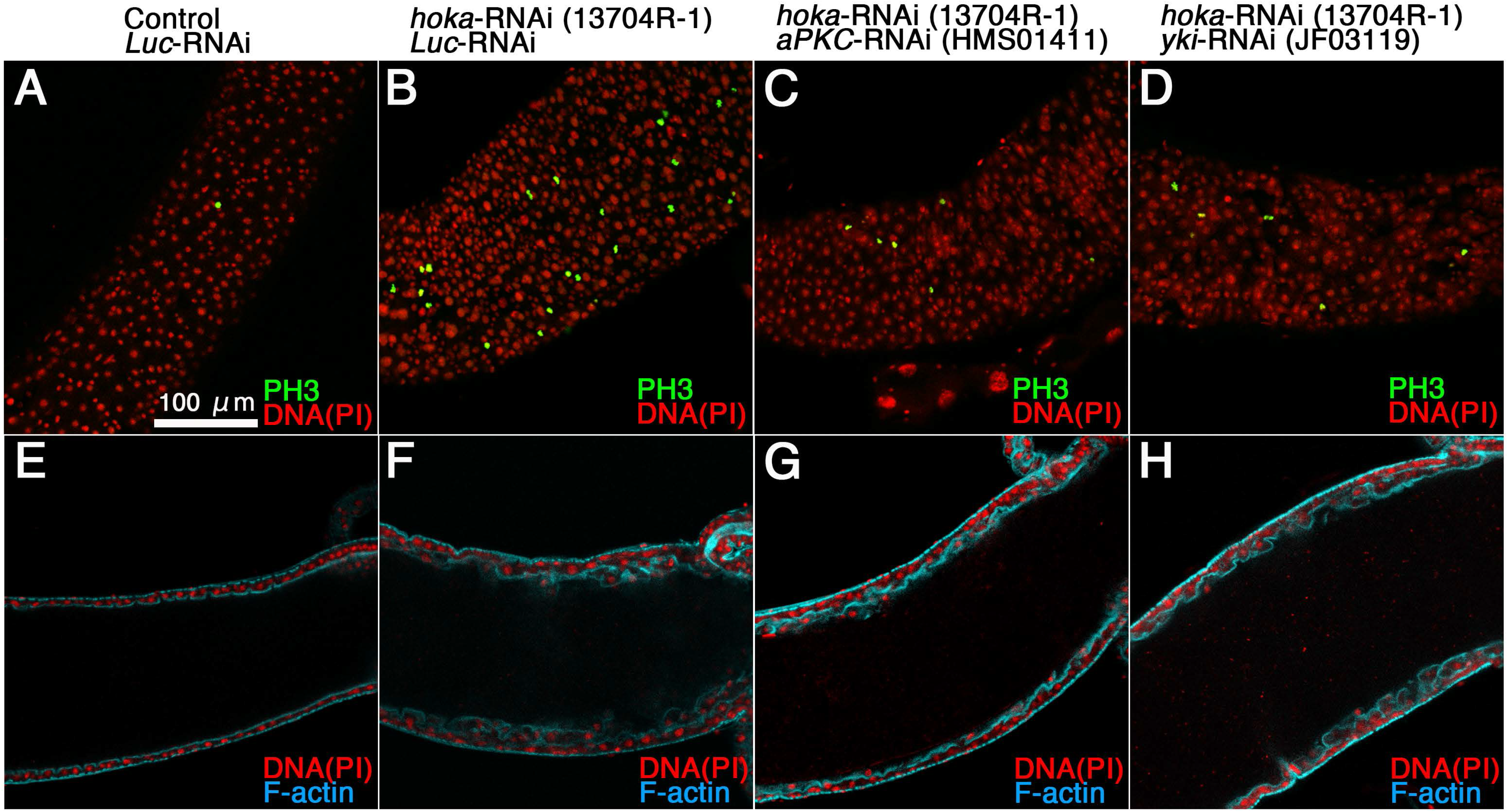
*aPKC*- and *yki*-RNAi together with *hoka*-RNAi in ECs results in reduced ISC overproliferation caused by *hoka*-RNAi. **(A–H)** Confocal images of the adult posterior midgut expressing *Myo1A^ts^-Gal4* with UAS-*Luc*-RNAi (control, A, E), UAS-*hoka*-RNAi 13704R-1 together with *Luc*-RNAi (B, F), UAS-*hoka*-RNAi 13704R-1 together with *aPKC-RNAi* HMS01411(C, G), or UAS-*hoka*-RNAi 13704R-1 together with *yki*-RNAi JF03119 (D, H) at 5 days after induction and stained for PH3 (green, A–D), DNA (propidium iodide, A–H) (red), and F-actin (E–H). The images (A–D) and (E–H) show the surface views of the midgut and longitudinal cross-sections through the center of the midgut, respectively. Scale bar (A–H): 100 μm.

